# Molecular basis for the host range function of the poxvirus PKR inhibitor E3

**DOI:** 10.1101/2024.05.16.594589

**Authors:** Sherry L. Haller, Chorong Park, Ryan C Bruneau, Dewi Megawati, Chi Zhang, Sameera Vipat, Chen Peng, Tatiana G. Senkevich, Greg Brennan, Loubna Tazi, Stefan Rothenburg

## Abstract

The antiviral protein kinase R (PKR) is activated by viral double-stranded RNA and phosphorylates translation initiation factor eIF2α, thereby inhibiting translation and virus replication. Most poxviruses contain two PKR inhibitors, called E3 and K3 in vaccinia virus (VACV), which are determinants of viral host range. The prevailing model for E3 function is that it inhibits PKR through the non-specific sequestration of double-stranded (ds) RNA. Our data revealed that Syrian hamster PKR was resistant to E3, which is at odds with the sequestration model. However, Syrian hamster PKR was still sensitive to K3 inhibition. In contrast, Armenian hamster PKR showed opposite sensitivities, being sensitive to E3 and resistant to K3 inhibition. Mutational analyses of hamster PKRs showed that sensitivity to E3 inhibition was largely determined by the region linking the dsRNA-binding domains and the kinase domain of PKR, whereas two amino acid residues in the kinase domain (helix αG) determined sensitivity to K3. Expression of PKRs in congenic cells showed that Syrian hamster PKR containing the two Armenian hamster PKR residues in helix-αG was resistant to wild type VACV infection, and that cells expressing either hamster PKR recapitulated the phenotypes observed in species-derived cell lines. The observed resistance of Syrian hamster PKR to E3 explains its host range function and challenges the paradigm that dsRNA-binding PKR inhibitors mainly act by the sequestration of dsRNA.

**Significance:** The molecular mechanisms that govern the host range of viruses are incompletely understood. A small number of poxvirus genes have been identified that influence the host range of poxviruses. We show that the host range functions of E3 and K3, two host range factors from vaccinia virus, are a result of species-specific interactions with the antiviral protein kinase R (PKR) and that PKR from closely related species displayed dramatic differences in their sensitivities to these viral inhibitors. While there is a substantial body of work demonstrating host-specific interactions with K3, the current model for E3-mediated PKR inhibition is that E3 non-specifically sequesters dsRNA to prevent PKR activation. This model does not predict species-specific sensitivity to E3; therefore, our data suggest that the current model is incomplete, and that dsRNA sequestration is not the primary mechanism for E3 activity.

## Introduction

Poxviruses exhibit wide variations in their host ranges, with some poxviruses such as variola virus, the causative agent of human smallpox, and swinepox infecting only a single host species. In contrast, other closely related poxviruses, such as cowpox viruses, monkeypox virus and vaccinia virus (VACV) infect and cause disease in many different mammalian species and are readily transmitted between different species (1, 2). Because poxviruses utilize receptors that are ubiquitous in most species, entry into host cells is largely species-independent (3). Instead, productive infection and therefore the host range of poxviruses is dependent on their ability to subvert the host’s innate immune responses (1). The molecular mechanisms underlying the differences in poxvirus host range are poorly understood. Through the use of deletion mutants and recombinant viruses, a number of viral genes that are necessary for replication in cells of some hosts but not in others have been identified and termed host range genes (4). Most of these poxvirus host range factors inhibit antiviral host proteins, including the protein kinase R (PKR) (1). PKR is an antiviral protein kinase, which is expressed at intermediate levels in most vertebrate cells and is up-regulated upon interferon stimulation. PKR encodes two dsRNA-binding domains in its N-terminus and a kinase domain in its C-terminus, which are connected by a linker region of variable length (5). During poxvirus infection, dsRNA is produced in the cytoplasm of infected host cells as a result of overlapping transcription of neighboring genes (6–8). PKR senses and binds this dsRNA, which results in a conformational change that allows two PKR monomers to dimerize and become autophosphorylated, which is necessary for PKR activation (9). Activated PKR then phosphorylates the alpha subunit of eIF2, which leads to a general shutdown of cap-dependent mRNA translation and inhibits viral replication in infected cells (5). PKR is a fast evolving gene, and the kinase domain of PKR has been evolving more rapidly than that of other eIF2α kinases, due to positive selection likely imposed by viral inhibitors (10, 11).

VACV, which is the prototypic member of the *Orthopoxvirus* genus, encodes two PKR inhibitors, called E3 (encoded by E3L, OPG65) and K3 (encoded by K3L, OPG41) (12, 13). Orthologs of E3L and K3L are found in the genomes of most chordopoxviruses (14). Like PKR, E3 is a dsRNA-binding protein and additionally contains a Z-DNA-binding domain (12, 15, 16). E3 is a multifunctional protein, which inhibits several cellular dsRNA-binding and Z-DNA-binding proteins in addition to PKR (17–19). However, in human HeLa cells, stable knock-down of PKR rescued replication of VACV lacking E3L, suggesting that PKR is the most important target of E3 in these cells (20). E3 prevents PKR autophosphorylation and therefore inhibits PKR at an early stage in the activation pathway. Therefore, the ability of E3 to bind and sequester viral dsRNA, shielding it from detection by cellular dsRNA-binding proteins such as PKR, is thought to be critical for its PKR-antagonizing activity (15, 21). This model is supported by experiments that replaced E3L with dsRNA-binding proteins from other viruses or even bacteria, which partially rescued replication of an E3L-deficient VACV in cell lines (22–26). In contrast, K3 inhibits PKR at a later stage after autophosphorylation. K3 contains an S1 domain with homology to the S1 domain of eIF2α and acts as a pseudosubstrate inhibitor of PKR, docking to the kinase domain of PKR and thus preventing interaction with and phosphorylation of eIF2α (27–30).

The host range functions of E3L and K3L were initially described in experiments with VACV strains in which either E3L or K3L was deleted. It was shown that E3L was required for VACV replication in human HeLa cells, but it was dispensable for virus replication in Syrian hamster BHK-21 cells. K3L was required for replication in BHK-21 cells but was dispensable for replication in HeLa cells. It was proposed that different levels of PKR expressed in each cell line and differences in the amount of dsRNA generated during infection might explain the differential requirement for E3L and K3L in these cells (21). In the time since these initial experiments, we and other labs have demonstrated that PKR from different species display differential sensitivity to K3 inhibition (10, 11, 30–33). Similarly, K3 orthologs from other poxviruses have been shown to inhibit PKR in a species-specific manner (31–36). To investigate whether E3 may also display species-specific PKR interactions, we used a sensitive cell-based luciferase assay to assess the sensitivity of PKR from different host species to inhibition by E3 or K3. We discovered substantial variations in the sensitivity of PKR from four related hamster species and identified two opposing instances of PKR resistance to inhibition by both viral proteins. Furthermore, the inability of E3 and K3 to inhibit PKR from these species correlated with replication of VACV mutants lacking E3L or K3L in cells from the corresponding hamster species, as well as with the phosphorylation of eIF2α by the endogenous PKR proteins. Using mutational analysis and domain swapping, we identified regions in hamster PKR orthologs that determine sensitivities to E3 or K3 inhibition. Together, these results support the conclusion that species-specific inhibition of PKR by VACV E3 and K3 contribute to their host range functions, and suggest that, at least in some cases the prevailing model of E3-mediated PKR inhibition by dsRNA sequestration is not the primary mechanism.

## Results

### Resistance of Syrian hamster PKR vaccinia virus E3

Previous results indicated that PKR from different mammalian species can be differentially inhibited by VACV K3 and K3 orthologs from other poxviruses in yeast-based growth assays and luciferase-based reporter (LBR) assays in PKR-deficient or depleted cells (10, 11, 30–33, 36), which explains the host range function of K3 orthologs. Intrigued by the finding that E3L but not K3L was dispensable for VACV replication in Syrian hamster BHK-21 cells (21), we hypothesized that Syrian hamster PKR might be resistant to E3 but sensitive to K3. We analyzed the sensitivity of PKR to E3 inhibition in the LBR assay by co-transfecting HeLa PKR^knock-down(kd)^ cells with either mouse or Syrian hamster PKR and increasing amounts of E3L. In this assay PKR gets likely activated by dsRNA that is formed by overlapping transcripts generated from the transfected plasmids (37). Whereas mouse PKR was inhibited by E3 in a dose-dependent manner, Syrian hamster PKR was resistant to E3 inhibition, even when high amounts of E3L were co-transfected (Fig. 1A). To determine if the phenomenon of resistance to E3 inhibition was unique to Syrian hamster PKR or whether PKR from other hamster species also share this trait, we cloned PKR from three additional hamster species: Turkish hamster (*Mesocricetus brandti*, Mb), Armenian hamster (*Cricetulus migratorius*, Cm) and Chinese hamster (*Cricetulus griseus*, Cg). Multiple sequence alignment and calculated protein identities show closer relatedness between Syrian and Turkish hamster PKR (97.3 % identity) and between Armenian and Chinese hamster PKR (89.9 % identity), whereas sequence identities between PKRs from the two *Mesocricetus* hamster species compared to PKR from the *Cricetulus* species ranged between 80.6 to 83% (Fig. S1A, B). We tested the four hamster PKRs as well as human and mouse PKR, for comparison, in the LBR assay for their sensitivities to VACV E3 and K3 (Fig. 1B, C). As expected, human and mouse PKR were both inhibited by E3. Both Armenian and Chinese hamster PKR were also sensitive to inhibition by E3. However, Turkish hamster PKR, like Syrian hamster PKR, was resistant to E3 inhibition (Fig. 1B). These results demonstrate that resistance to E3 is not unique to Syrian hamster PKR but is shared by at least one other *Mesocricetus* species’ PKR.

**Fig. 1.**
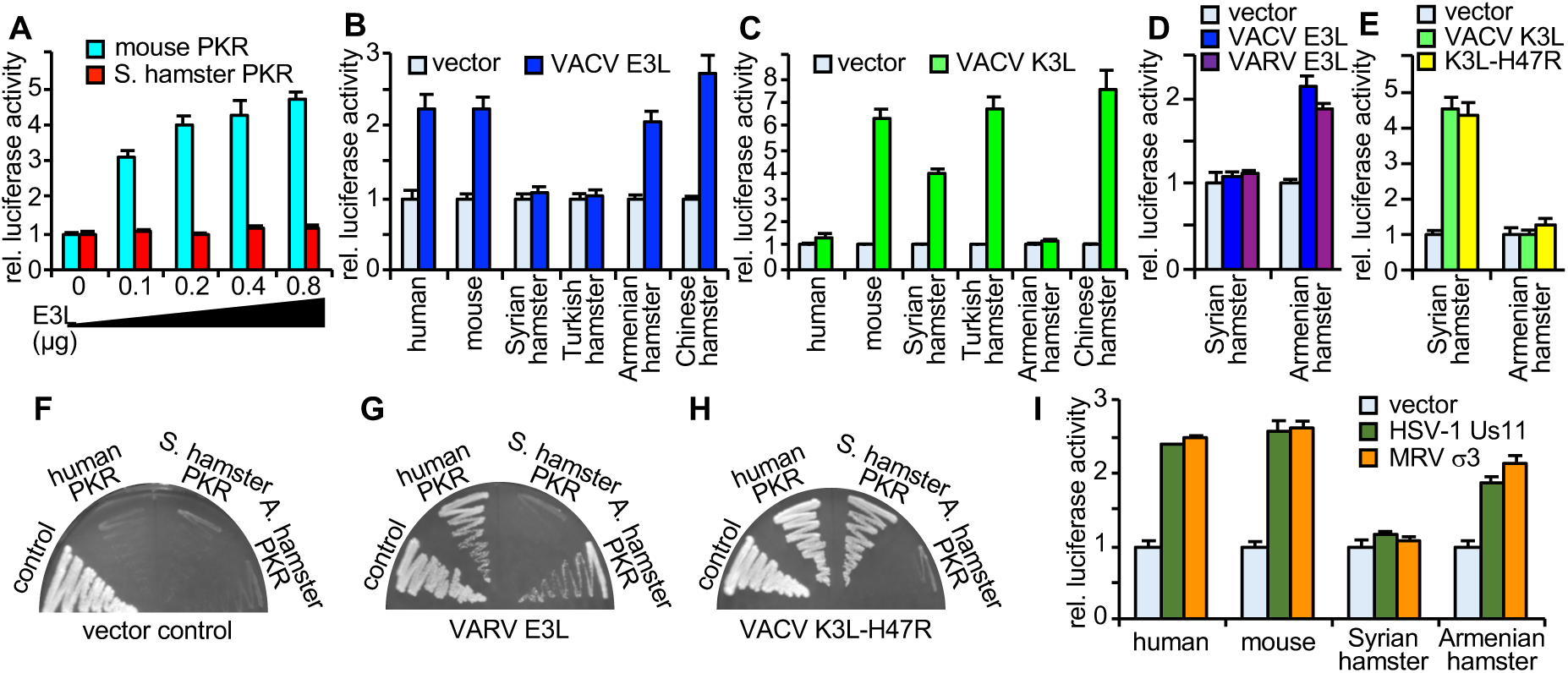
Species-specific inhibition of PKR by vaccinia virus E3 and K3. HeLa-PKR^kd^ cells were transfected with plasmids encoding firefly luciferase (0.05 µg), the indicated PKRs (0.2 µg), and the indicated inhibitors (0.4 µg, or as indicated in (A)). Luciferase activities were determined 42-48 hours after transfection and results were normalized to PKR-only transfected cells. Error bars indicate the standard deviations for three replicate transfections. (A) Plasmids encoding Syrian hamster or mouse PKRs were co-transfected with increasing amounts of VACV E3L. (B, C) PKR from the indicated species were co-transfected with E3L or K3L. (D, E) Syrian or Armenian hamster PKR were co-transfected with VACV E3L or VARV E3L (D), or VACV K3L or VACV K3L-H47R (E). (F-H) Plasmids encoding human, Syrian or Armenian hamster PKR under the control of a yeast GAL-CYC1 hybrid promoter were transformed into yeast strains that were stably transformed with the empty vector (control) (F), VARV E3L (G), or VACV K3L-H47R (H). Transformants were colony-purified under non-inducing conditions and then streaked on SC-Gal plates to induce PKR expression for 7 days. Representative results are shown for 4 independent transformations. (I) Plasmids encoding the indicated PKRs were co-transfected with HSV-1 Us11, or mammalian reovirus (MRV) σ3.

When we assessed K3 sensitivity in this assay, human PKR, as expected, was largely resistant to inhibition by K3, whereas mouse, Syrian hamster and Turkish hamster PKR were efficiently inhibited. However, PKR from the *Cricetulus* hamster species exhibited different sensitivities to K3. Chinese hamster PKR was inhibited by K3, whereas Armenian hamster PKR was resistant (Fig. 1C). In order to compare expression levels of Syrian and Armenian hamster PKR, we transfected HeLa PKR^kd^ cells with FLAG-tagged PKR, either alone or in combination with FLAG-tagged K3 or E3 and performed western blot analyses with cell lysates (Fig. S2). Both hamster PKR were expressed at comparable levels in the absence of inhibitors. Whereas co-transfection with an ineffective inhibitor did not change PKR expression levels, co-transfection with an effective inhibitor led to increased PKR expression, as expected because autoinhibition of PKR expression was derepressed. We also tested sensitivity patterns of Syrian hamster and Armenian hamster PKR to the E3 ortholog from variola virus (VARV) and a variant of VACV K3 (H47R), which was identified as a better inhibitor of human PKR than wild type K3 in a yeast assay (38). Syrian hamster PKR was resistant to both E3 orthologs, whereas Armenian hamster PKR was sensitive to both (Fig. 1D). Likewise, K3 and K3L-H4R showed comparable inhibition of Syrian hamster PKR (sensitive) and Armenian hamster PKR (resistant) (Fig. 1E).

In complementary experiments, we tested the sensitivities of Syrian and Armenian hamster PKR to E3 and K3 inhibition using a yeast-based growth assay involving two previously described yeast strains that inducibly express either VARV E3 or a variant of VACV K3-H47R. Plasmids encoding human, Syrian hamster, or Armenian hamster PKR, as well as an empty control plasmid, which are all under the control of a yeast GAL-CYC1 hybrid promoter, were transformed into yeast strains stably expressing either the empty vector, VARV E3 or VACV K3-H47R. In the control strain, induction of all PKRs prevented yeast growth compared to yeast transformed with the empty vector (Fig. 1F). The toxicity of human and Armenian hamster PKR, but not of Syrian hamster PKR, was suppressed when VARV E3 was co-expressed (Fig. 1G). In VACV K3-H47R-expressing cells toxicity of human and Syrian hamster PKR was suppressed, whereas Armenian hamster PKR still suppressed yeast growth. The results of these yeast assays corroborate the results obtained in the LBR assays and demonstrate differential sensitivity of Syrian and Armenian hamster PKR to E3 and K3 inhibition.

In order to analyze whether the resistance of Syrian hamster PKR also extends to dsRNA-binding PKR inhibitors from other viruses, we co-transfected PKR with the previously characterized PKR inhibitors Us11 from Herpes-simplex virus 1 (HSV-1) (39, 40) and σ3 from mammalian reovirus (41, 42). Human, mouse, and Armenian hamster PKR were efficiently inhibited by Us11 or σ3. In contrast, Syrian hamster PKR was resistant to both Us11 and σ3 (Fig. 1I). These results show that Syrian hamster PKR, unlike other tested PKRs, is resistant to multiple, unrelated dsRNA-binding inhibitors from a variety of viruses.

### Contrasting VACV replication in cell lines from different hamster species

It is noteworthy that Syrian hamster BHK-21 is the only hamster cell line in which VACV is routinely propagated, and that VACV is unable to replicate in Chinese hamster CHO-K1 cells due to the induction of cell death (43). In order to evaluate if the species-dependent inhibition of hamster PKR that we observed in the LBR and yeast assays correlates with VACV infection, we first tested whether VACV was able to replicate in cells from different hamster species. We infected five hamster cell lines, derived from three different species, at a multiplicity of infection of 0.01 to observe multi-cycle replication with wild type VACV-Copenhagen (VC-2) and compared these titers to a derived VACV strain in which both E3L and K3L were deleted (VC-R2, for simplicity referred to as VC-ΔE3ΔK3) (Fig. S3). BHK-21 cells derived from a Syrian hamster and AHL-1 cells derived from an Armenian hamster were both permissive to VACV infection, whereas VC-ΔE3ΔK3 was unable to replicate above input levels in either cell line. In agreement with the previous reports, neither wild type VC-2 nor VC-ΔE3ΔK3 could replicate in CHO-K1 cells. Because it was not clear if the inability of VACV to replicate in CHO cells was due to cell-type-specific or a species-specific effect, we tested two additional Chinese hamster cells lines, Don and V79-4, for their ability to support VACV replication. VC-2 replicated to titers about 100-400-fold above input in both additional Chinese hamster cell lines, although the titers were lower than those in BHK-21 and AHL-1 cells. VC-ΔE3ΔK3 did not replicate in either Don or V79-4 cells. We decided to use V79-4 Chinese hamster cells in subsequent experiments because their cell-doubling time is more similar to the BHK-21 and AHL-1 cells, whereas the Don cells grow more slowly.

### eIF2α phosphorylation and VACV replication in hamster cells correlates with PKR sensitivity to E3 and K3

When the host range functions of E3 and K3 were initially described, either E3L or K3L-deficient VACV strains were compared to the parental virus. To extend this analysis, we compared virus replication of wild type VC-2, VC-R1 (ΔE3L, for simplicity referred to as VC-ΔE3, vP872 (ΔK3L, for simplicity referred to as VC-ΔK3), and VC-ΔE3ΔK3. In HeLa-PKR^kd^ cells all viruses were able to replicate, including VC-ΔE3ΔK3, which showed about 5,000-fold increased replication in comparison to HeLa control cells (Fig. 2C). We used these viruses to determine whether E3 or K3 may be more important for VACV replication in certain hamster cell lines by measuring eIF2α phosphorylation. We infected BHK-21, AHL-1, and V79-4 cells with the four virus strains for 6 hours (MOI = 5.0) and performed western blot analyses. In all three cell lines, low levels of eIF2α-P were observed in uninfected and VC-2 infected cells, whereas high levels were detected after infection with VC-ΔE3ΔK3 (Fig. 2A, B). In BHK-21 cells, infection with VC-ΔE3 did not result in increased eIF2α phosphorylation, whereas infection with VC-ΔK3 resulted in comparable eIF2α phosphorylation as VC-ΔE3ΔK3. The opposite phenotypes were observed in AHL-1 cells, in which infection with VC-ΔE3 but not VC-ΔK3 resulted in strong eIF2α phosphorylation. In V79-4 cells, infection with both VC-ΔE3 and VC-ΔK3 resulted in less eIF2α phosphorylation than VC-ΔE3ΔK3. These data show that K3L was required to suppress eIF2α phosphorylation in BHK-21 cells and that E3L was required to suppress eIF2α phosphorylation in AHL-1 cells.

**Fig. 2.**
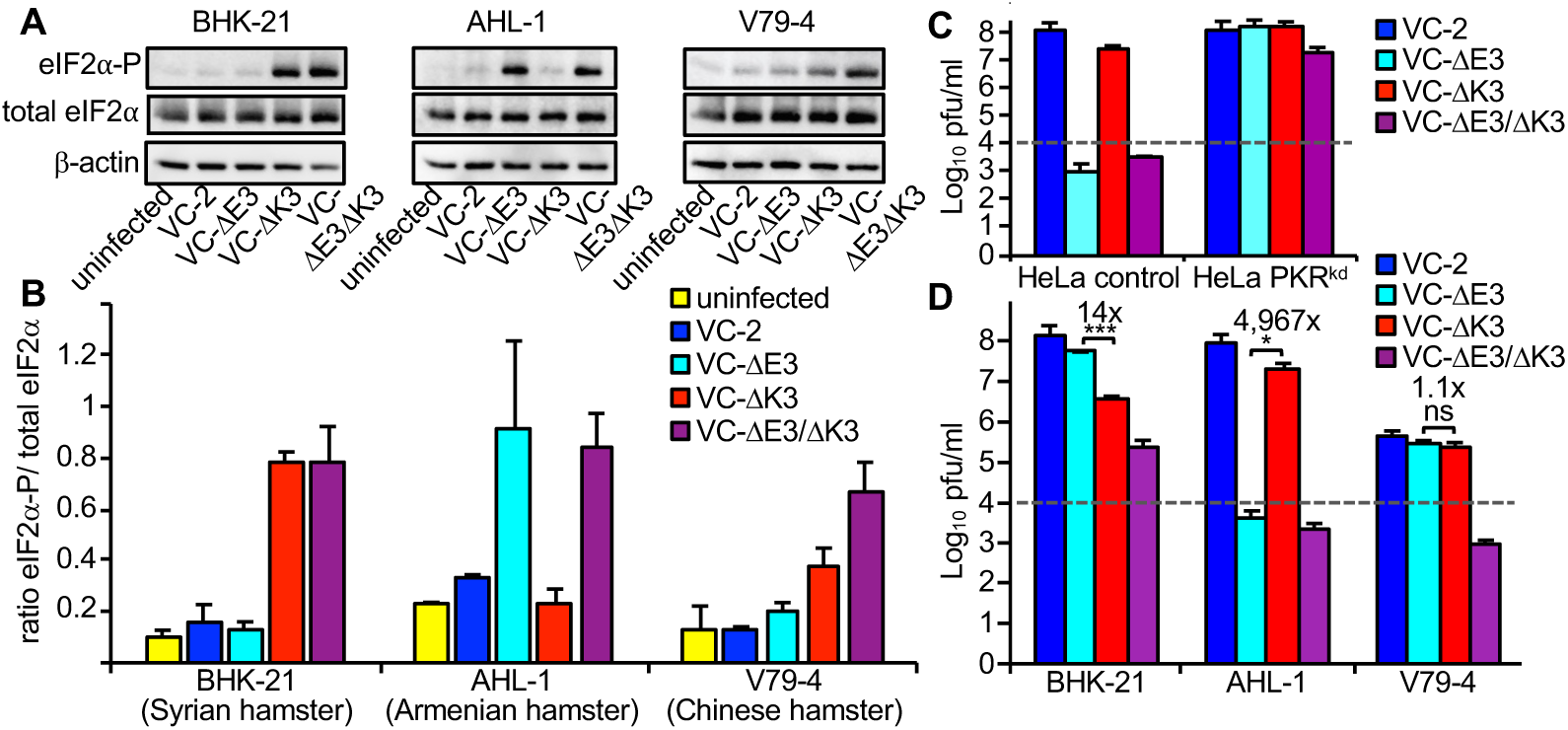
Effects of E3 and K3 on eIF2a phosphorylation and VACV replication. (A) Three hamster cell lines were infected (MOI = 5) with wild type VACV-Cop (VC-2) or derived mutant viruses lacking E3L (VC-R1), K3L (vP872), or both (VC-R2) at MOI=5. Cellular lysates were collected 6 hpi and analyzed by western blot for phosphorylated eIF2a (Ser-51, top panels), total eIF2a (middle panels), or b-actin (bottom panels). (B) Band intensities were measured using the Kodak-400MM Image station software and the ratios of phosphorylated eIF2a to total eIF2a were calculated and plotted on the graph. Error bars indicate the standard deviations of two independent experiments. (C) HeLa control and HeLa PKR^kd^ cells were infected with the indicated VACV strains (MOI = 0.01) for 30 h and titered on RK13+E3L+K3L cells. (D) Hamster cell lines were infected with wild type VACV strains (MOI = 0.01) for 30 h and titered on RK13+E3L+K3L cells. Error bars indicate the standard deviation of two experiments. Fold differences between the single-deletion mutant viruses are noted above each pair. P-values were calculated using the Student’s t-test (* = p < 0.05; *** p = < 0.0005).

To analyze the effects of E3L and K3L on virus replication, the same hamster cell lines were infected with all four VACV strains (MOI = 0.01). After 48 hours, virus supernatants were collected and titered on RK13+E3L+K3L cells. In BHK-21 cells, VC-ΔE3ΔK3 showed weak replication (∼ 20-fold increase over input), whereas no replication was observed in AHL-1 and V79-4 cells (Fig. 2D). VC-ΔE3 and VC-ΔK3 showed opposing replication profiles in BHK-21 and AHL-1 cells: In BHK-21 cells, VC-ΔE3 replicated significantly better (about 10-fold) than VC-ΔK3, whereas in AHL-1 cells VC-ΔK3 replicated to much higher titers than VC-ΔE3. Both VC-ΔE3 and VC-ΔK3 replicated to comparable levels in V79-4 cells and slightly less than VC-2 (∼2-5-fold reduction in repeated experiments), suggesting that PKR in these cells was susceptible to inhibition by either E3 or K3 and that these viral inhibitors might have additive effects. Overall, the observed differences in the replication assays correlated well with the susceptibility of PKR from the respective hamster species observed in the other assays.

### Interferon-induced PKR expression augmented the replication differences between K3L and E3L-deficient VACV in BHK-21 cells

The difference between the replication of E3L- and K3L-deficient strains observed in our experiments were not as pronounced as the differences previously reported in BHK cells (21). To analyze whether PKR can be induced in the hamster cell lines, and whether increased PKR expression may enhance the differences we observed, we incubated hamster cells with mouse interferon (IFN)-α1 for 17 h and analyzed mRNA expression using semi-quantitative RT-PCR. IFN-α1 treatment induced PKR expression in BHK-21 and AHL-1 cells, but not in V79-4 cells (Fig. 3A, B). We next treated the same cell lines with IFN-α1 for 24 hours before infecting them with the VACV strains. IFN-α1 treatment of BHK-21 cells reduced VC-2 and VC-ΔE3 replication approximately 10-20-fold, but we observed stronger reduction of VC-ΔK3 replication (∼200-fold), relative to untreated cells (Fig. 3B). The differences in replication between VC-ΔE3 and VC-ΔK3 were approximately 10-fold increased after IFN-treatment. In AHL-1 cells, IFN-α1 treatment reduced virus replication for all tested strains, but the relative differences between strains remained comparable. No substantial differences in virus replication were observed in V79-4 cells, consistent with the lack of IFN-mediated induction of PKR.

**Fig. 3.**
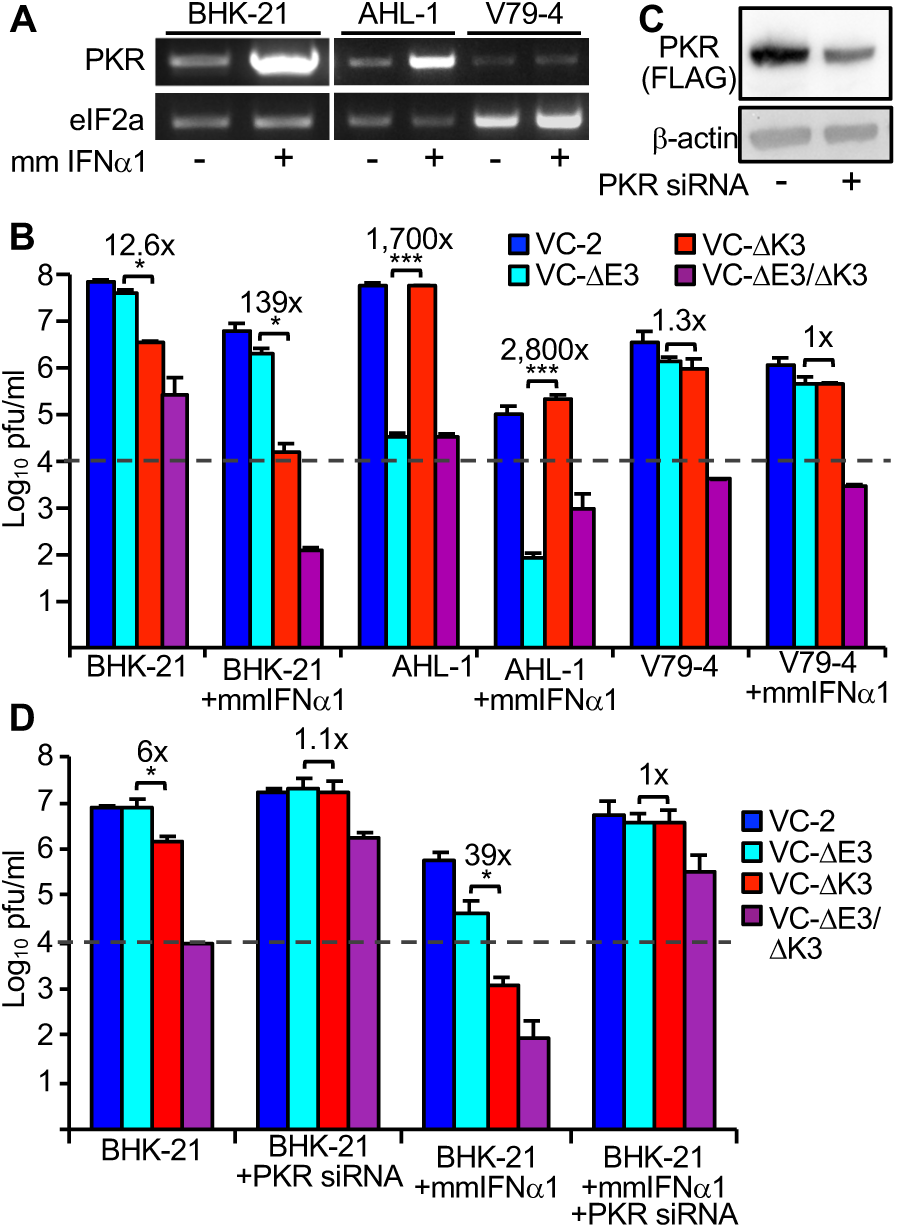
Effect of interferon treatment on VACV replication in hamster cells. (A) Total RNA was isolated from hamster cells either untreated (-) or treated (+) with 5 units/ml of mouse (mm) interferon (IFN)*α*1 for 17 hours. PKR and eIF2a genes were amplified by RT-PCR. (B) Hamster cell lines were left untreated or treated with 50 units/ml mmIFN*α*1 for 24 hours, and then infected with the indicated VACV strains (MOI = 0.01). Lysates were collected at 30 hpi and titered. Fold differences between the single deletion mutant viruses are indicated above each pair. Error bars indicate the standard deviation of two experiments. P-values were calculated using the Student’s t-test (* = p < 0.05, ***p = < 0.0005). (C) T-REx 293 PKR^ko^ + Syrian hamster PKR cells were induced with doxycycline 24 h prior to treatment with pooled siRNA targeting hamster PKR. Cell lysates were collected after 24 h and analyzed by western blot to detect FLAG-tagged PKR and b-actin. (D) BHK-21 cells were mock-transfected or transfected with pooled siRNA targeting hamster PKR and were either left untreated or treated with mmIFN*α*1 (50 U/ml). 24 h after treatment, cells were infected with the indicated VACV strains (MOI = 0.01). Lysates were collected 30 hpi and titered. Fold differences between the single deletion mutant viruses are noted above each pair. Error bars indicate the standard deviation of three replicate experiments. P-values were calculated using the Student’s t-test (* p = < 0.05).

In order to investigate whether knock-down of PKR differentially affects replication of the VACV strains, we treated BHK cells with hamster PKR-specific siRNA. No antibodies against Syrian hamster PKR are commercially available. As an alternative approach, we used PKR knock-out 293 T-REx cells to express FLAG-tagged Syrian hamster PKR (described below). Treatment with the siRNA substantially reduced expression of PKR (Fig. 3C). Knock-down of PKR resulted in comparable replication of VC-2, VC-ΔE3 and VC-ΔK3 in BHK cells, and ∼200-fold increased replication of VC-ΔE3ΔK3, although, likely due to the incomplete knock-down, not to the same level as observed with the other strains (Fig. 3D). PKR knock-down also reversed the effects of IFN-α1 treatment on the replication of all four strains indicating that the observed antiviral effects of IFN-α1 were largely mediated by PKR.

### Identification of regions in PKR determining sensitivity to inhibition by E3 and K3

The region linking the dsRNA-binding domains with the kinase domains is highly divergent between PKRs from different species in respect to amino acid composition and length. Thus, we hypothesized that this region may influence PKR sensitivity to E3. We tested a naturally occurring splice variant (sv) of Syrian hamster PKR, in which 54 out of 85 amino acids comprising the linker are missing (Fig. S1). This splice variant was as resistant to E3 inhibition as full-length PKR (Fig. 4A). We subsequently removed more parts of the linker region in blocks of six additional amino acids per construct (Fig. S1). Removal of 12 additional amino acids (Δ164-229) increased the sensitivity of this PKR to E3 inhibition, implicating a role for the linker region in resistance to E3 (Fig. 4A). We next constructed hybrids between Syrian and Armenian hamster PKR to independently determine the regions involved in E3 and K3 sensitivity (Fig. 4B). Swapping the kinase domains between the PKRs showed that the sensitivity to K3 was determined by the kinase domain, whereas sensitivity to E3 was determined by a region comprising the dsRBDs and the linker (Fig. 4C). Swapping only the first dsRBD1 did not affect sensitivity as compared to wild type PKRs. However, including the second dsRBD2 resulted in intermediate sensitivity of the hybrids, whereas addition of the linker region was necessary to phenocopy sensitivity of the wild type PKRs (Fig. 4D). In another set of experiments, we swapped only individual regions dsRBD1, dsRBD2, or the linker region between the two PKRs. Here, only hybrids with the swapped linker region showed completely reversed sensitivity to E3 inhibition (Fig. 4E). These results show that the linker region is the primary determinant of hamster PKR sensitivity to E3 inhibition, although dsRBD2 might play a secondary role.

**Fig. 4.**
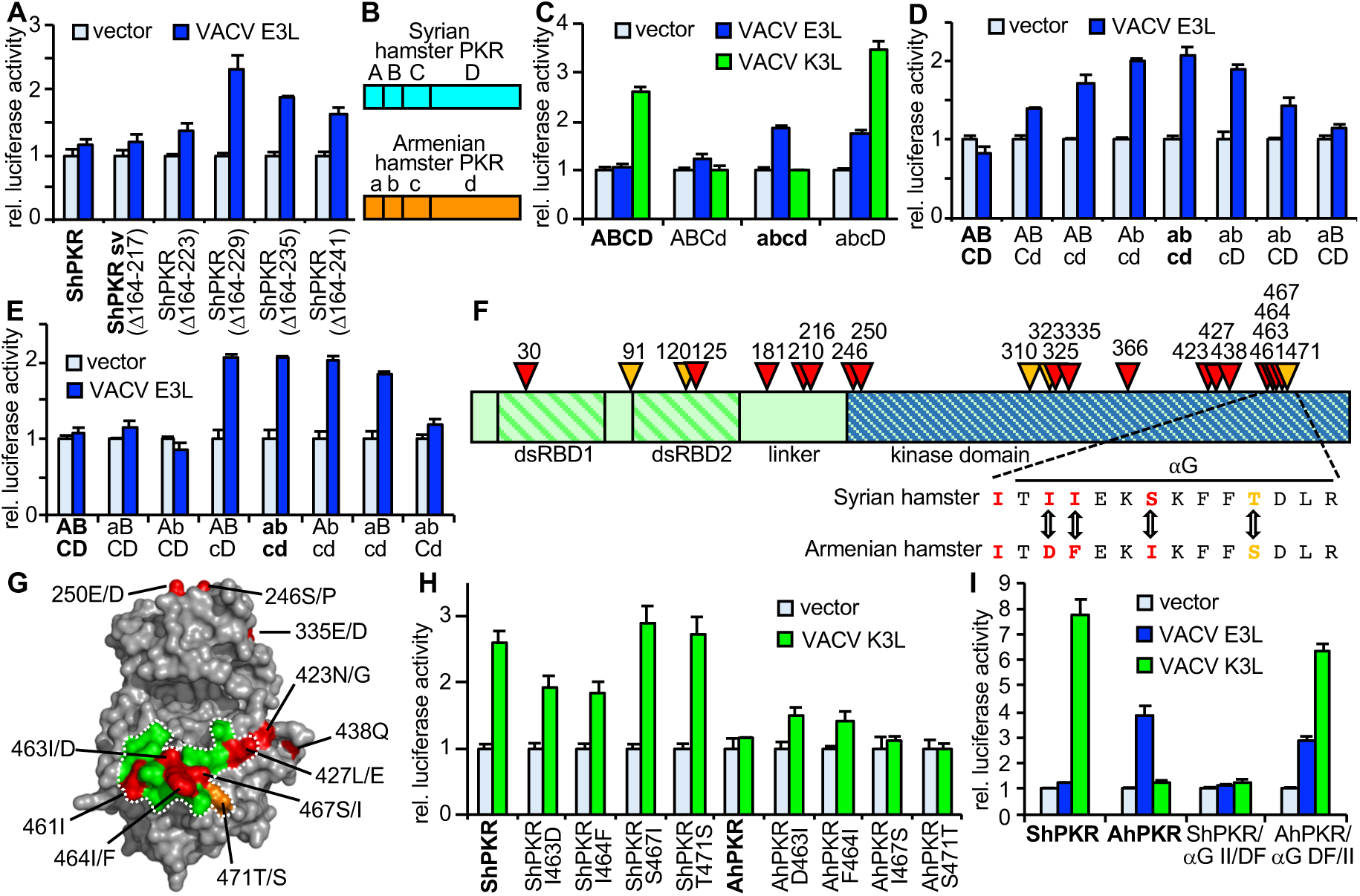
Defining the regions of hamster PKRs conferring sensitivity to E3 and K3. HeLa-PKR^kd^ cells were co-transfected with plasmids encoding luciferase (0.05 µg), PKR, or PKR mutants (0.2 µg) and VACV E3L or VACV K3L (0.4 µg). Luciferase activities were determined 42-48 hours after transfection. For each transfection luciferase activity was normalized to PKR-only transfected cells. Error bars indicate the standard deviations for three replicate transfections. Wild type PKRs are indicated with bold letters. (A) Syrian hamster PKR, a splice variant (sv), or mutants in which blocks of 6 additional amino acids were removed from PKR sv were tested for sensitivity to E3. (B) Schematic of the domain structure used for chimeric PKR construction. Letters denote different PKR domains: A/a = first dsRBD, B/b = second RBD, C/c = linker region, D/d = kinase domain. (C-E) Sensitivities of chimeric PKR to E3 and K3. Wild type PKRs are indicated with bold letters. (F) Positively selected residues in *Cricetidae* and *Muridae* PKRs were determined using PAML and are projected on the domain structure of Syrian hamster PKR. Red and orange triangles indicate positions of positively selected residues (p ≥ 0.95) that were detected in both datasets, or in one of the datasets, respectively. (G) Positively selected residues projected on the crystal structure of human PKR (44). Residues are numbered in respect to Syrian hamster PKR. Residues after slashes indicate the Armenian hamster residue if they are divergent. Dotted line encompasses PKR residues that contacted eIF2a in the co-crystal-structure (44). (H) Positively selected residues in helix αG of Syrian and Armenian hamster PKR were swapped as indicated, and sensitivities to K3 inhibition were assessed. (I) Sensitivity of mutant hamster PKRs, in which both residues 463 and 464 were swapped between Syrian and Armenian hamster PKR, to E3 and K3 inhibition.

Key residues governing host-virus interactions are often subjected to strong selective pressure from both host and virus and often exhibit signatures of positive selection. To identify positively selected residues in hamster and hamster-related PKRs, we gathered high-quality PKR sequences from 36 rodent species from GenBank and performed a phylogenetic analysis (Fig. S4 and Table S1). We next performed PAML analysis with all available sequences from the families *Cricetidae* and *Muridae*, with or without sequences from the *Neotominae* subfamily, because the sequences from the latter contain shorter linker regions. 16 residues were identified as positively selected (P ≥ 0.95%) in both datasets with model 8 (indicated by red triangles in Fig. 4G, and Table S2), with six additional residues that were significant in only one of the two datasets (orange triangles). Projection of the residues on the crystal structure of the human PKR kinase domain (44), revealed clustering of positively selected residues around the eIF2α-binding site of PKR, including four residues in helix αG that are divergent between Syrian and Armenian hamster PKR (Fig. 4F, G). These results were consistent with our previous experiments, which demonstrated that the kinase domain primarily influences K3 sensitivity in other species. To test this hypothesis, we first swapped the residues individually between the two hamster PKRs and determined sensitivity to K3. Mutation of residues 463 and 464 of Syrian hamster PKR resulted in slightly less sensitivity to K3 inhibition, whereas the corresponding mutations slightly increased the sensitivity of Armenian hamster PKR. Mutation of residues 467 or 471 had no effects on sensitivity to K3 (Fig. 4H). Mutation of both residues 463 and 464 (PKR-αG II/DF or PKR-αG DF/II) reversed this sensitivity making Syrian hamster PKR resistant and Armenian hamster PKR sensitive to K3 inhibition. These mutations did not alter the sensitivity to E3 (Fig. 4I). Syrian hamster PKR-αG II/DF was thus resistant to both E3 and K3.

### Syrian hamster PKR helix αG mutant provides strong antiviral activity against wild type vaccinia virus

To investigate PKRs from different species in a congenic background, we first knocked-out endogenous PKR in Flp-In T-REx 293 (T-REx) cells using CRISPR-Cas9. These cells contain a single insertion site for the transgene which is controlled by a tetracycline-inducible promoter. We selected clone A2, which was permissive for the PKR-sensitive VC-ΔE3ΔK3 for further characterization (Fig. 5A,B). In this clone, we did not detect PKR by western blot analysis (Fig. 5C). To confirm the knock-out we sequenced both genomic DNA and cDNA and identified a single 17 bp deletion, which overlaps with the guide RNA target site and no evidence of wild type PKR sequence (Fig. S5). We next stably inserted FLAG-epitope-tagged Syrian hamster PKR, Armenian hamster PKR, Syrian hamster PKR-αG II/DF, or human PKR into the FRT site in the T-REx PKR^ko^ cells. In all cells, doxycycline treatment induced PKR expression, although Armenian hamster PKR was expressed slightly less than the other PKRs (Fig. 5D). Furthermore, all of these PKR-expressing cells strongly suppressed VC-ΔE3ΔK3 replication, showing that PKRs are functional and that replication of VC-ΔE3ΔK3 in the PKR^ko^ cells is due to the lack of PKR (Fig. 5E). Cells expressing Syrian hamster PKR showed no suppression of the K3L-containing viruses and about 50-fold reduction of VC-ΔK3 replication. Armenian hamster PKR expression resulted in no suppression of the E3L-containing viruses, but strong repression of VC-ΔE3 replication. These results thus mimic those obtained with the Syrian hamster BHK and Armenian hamster AHL-1 cells. Infection of the Syrian hamster PKR-αG II/DF-expressing cells showed strong suppression of all tested viruses, including wild type VC-2. These results demonstrate a causal connection between PKR sensitivity to E3 and K3 and inhibition of virus replication and indicate that recombinant PKRs can substantially suppress the replication of a wild type poxvirus.

**Fig. 5.**
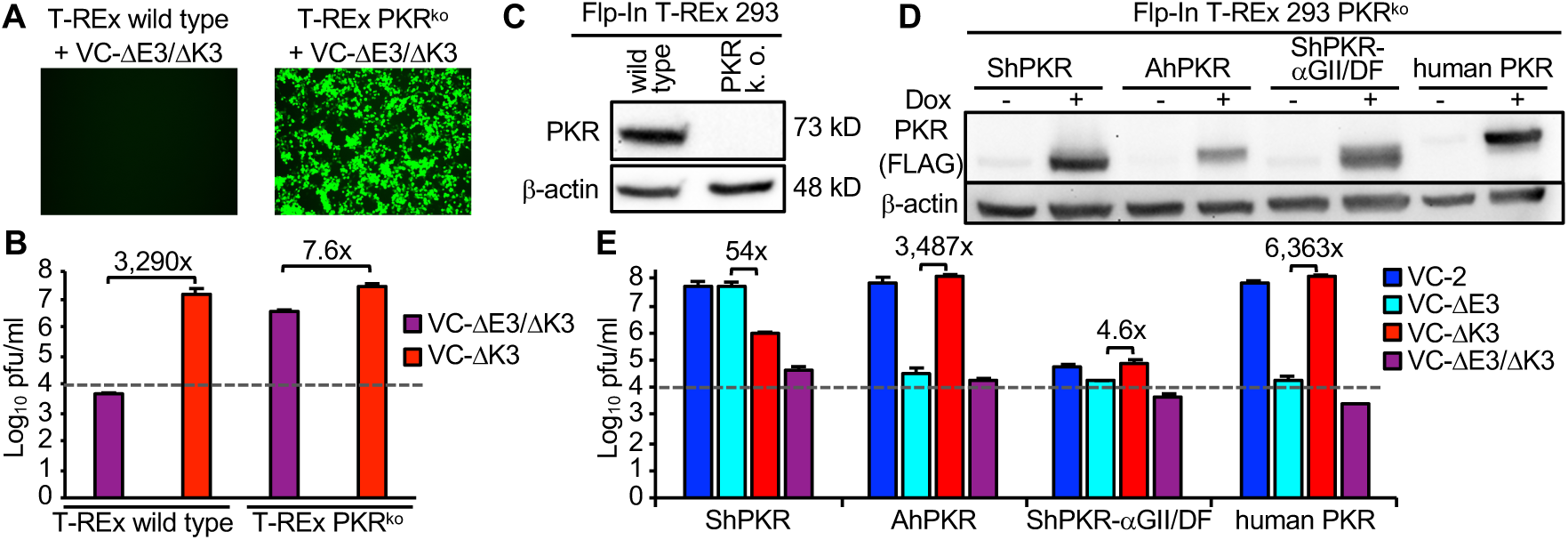
Generation of congenic PKR cells and effect on VACV replication. (A) Wild type and PKR^ko^ T-Rex 293 cells were infected with VC-R4 at MOI = 0.01. 45 hours after infection cells were visualized by fluorescence microscopy (magnification: 50x; exposure time: 150ms). (B) Parental and PKR^ko^ T-REx cells were infected in triplicates with VC-R4 or vP872 at MOI = 0.01 for 45 hours. Viral titers were determined by plaque assay. (C) Western blot of cell lysates from parental and PKR^ko^ T-REx cells using anti-PKR and anti-β-actin antibodies. (D) Expression of PKRs in T-REx PKR^ko^ cells. PKR expression in the indicated congenic cell lines was induced for 24 hours by doxycycline, or cells were left untreated. Total protein lysates were analyzed by western blot using anti-FLAG and anti-β-actin antibodies. (E) PKR-expression in congenic cells was induced for 24 hours before infection with the indicated VACV strains for 48 hours at MOI = 0.01. Error bars indicate the standard deviation of two experiments and fold differences between the single-deletion mutant viruses are noted above each pair.

## Discussion

Poxvirus host range genes are only required for productive replication in a subset of host cells and species (1, 4). While many poxvirus host range factors are immunomodulators, the molecular mechanisms underlying their host range function is, in most cases, poorly understood. Here we provide molecular explanations for the host range functions of VACV E3L and K3L in cells from different hamster species.

In this study we observed three different patterns for PKR susceptibility to E3 and K3 inhibition: (i) E3L but not K3L was dispensable for VACV replication in Syrian hamster BHK-21 cells, as previously reported (21); (ii) in Armenian hamster cells, K3L but not E3L was dispensable for virus replication; (iii) in Chinese hamster V79-4 cells, either E3L or K3L was dispensable for virus replication. VACV replication correlated well with eIF2α phosphorylation levels during infection in these cell lines, and with the sensitivities of the respective PKRs to E3 and K3 inhibition in reporter assays. These correlations were also consistent in infections of congenic cells expressing either Syrian or Armenian hamster PKR, which supports a causal connection between the presence of E3L or K3L, PKR and virus replication. Because many poxviruses possess two PKR inhibitors, this redundancy allows virus replication in cases in which a given PKR is resistant to one inhibitor, e.g. Syrian and Armenian hamster PKR. In cases in which PKR is sensitive to both inhibitors, e.g. Chinese hamster PKR, the loss of one inhibitor can be compensated by the other. In this system, a condition of being recognized as a host range factor is that PKR from a given species must have opposing sensitivities to one PKR inhibitor. This study therefore defines substantial variability in the redundancy of these two distinct host range factors in multiple hamster species/cells.

The resistance of Syrian and Turkish hamster PKR to E3 inhibition was unexpected and is at odds with the prevailing model of how E3 and other viral dsRNA-binding proteins inhibit PKR. According to this sequestration model, E3 sequesters viral dsRNA as a function of its dsRNA binding properties, thereby preventing detection by PKR and other dsRNA binding proteins (15, 21). Under this model, dsRNA sequestration would be predicted to inhibit PKR from all species; therefore, our discovery of hamster PKRs differentially susceptible to E3 cannot be explained by this model. Previous studies showed that E3 mutants with a reduced ability to bind dsRNA were still able to inhibit human PKR (45), and that E3 orthologs from sheeppox virus and Yaba monkey tumor virus were inefficient inhibitors of PKR despite dsRNA-binding capabilities (31, 46), which indicated that the sequestration model does not fully explain E3 activity. An alternative model for how E3 inhibits PKR activation proposes that E3 prevents PKR homodimerization, an essential step during PKR activation, by forming heteromers with PKR (47, 48). Supporting this heterodimer model, the myxoma virus E3 ortholog M029 was shown to interact with rabbit PKR in a dsRNA-dependent manner (49). This model is compatible with the mechanisms used by other viruses, for example viral RNA inhibitors VA(I) from human adenovirus and EBER(I) from Epstein-Barr virus, which inhibited PKR activation by preventing PKR homodimerization (50). In light of these data, one explanation for the resistance of Syrian and Turkish hamster PKR is that these host proteins might resist the formation of heterodimers with E3 and could therefore still form active homodimers.

Syrian hamster PKR showed complete resistance to E3 in the LBR assay, and the presence of E3 alone during VACV infection did not reduce eIF2α phosphorylation levels in BHK-21 cells. However, VACV with E3 as the only PKR inhibitor replicated about 10-fold better than VACV missing both E3 and K3. These data indicate a minor role for E3 in VACV replication in BHK-21 cells. PKR knock-down abolished these differences in replication between E3- and K3-only -expressing viruses, supporting the idea that this is a PKR-dependent phenotype. The relatively weak effect of E3 in these cells, may be due to some dsRNA sequestration, even if this this effect could not be readily measured by eIF2α phosphorylation levels. This sequestration effect might be more relevant in cells expressing relatively low levels of PKR, as induction of PKR expression by IFN increased the replication differences between K3-, and E3-only expressing viruses, an effect that was also abolished by PKR-knock-down. Low PKR levels in BHK-21 cells could also explain why VACV lacking both decapping enzymes D9 and D10, which results in elevated dsRNA levels, was able to replicate in BHK-21 cells (51). In summary, neither the dsRNA sequestration model nor the E3/PKR heteromer model on their own can fully explain the results presented here, indicating that both mechanisms may contribute to PKR inhibition by E3, with dsRNA sequestration by E3 playing a subordinate role to E3’s prevention of PKR homodimerization.

Domain swapping between Syrian and Armenian hamster PKR revealed that the sensitivity to E3 inhibition was largely determined by the region linking dsRBD2 with the kinase domain, and potentially a minor contribution from dsRBD2. This was supported by the finding that partial deletion of the linker in Syrian hamster PKR resulted in sensitivity to E3 inhibition. The linker region is highly basic and has been described as unstructured and highly flexible in solution (52, 53). The linker has also been implicated in inhibiting PKR activation in its non-dsRNA bound inactive state (9, 54). During PKR activation, the very N-terminal residues of the kinase domain were shown to be perturbed indicating that the conformation of the linker changes during this process (52). Taken together, these studies strongly suggest that the linker has an important and relatively unrecognized role in PKR activation and evasion of viral antagonists.

The identification of amino acid residues and regions in Syrian and Armenian hamster PKR that determine sensitivity to K3 and E3 allowed us to generate recombinant PKR that was resistant to both E3 and K3. Importantly, expression of this E3/K3-resistant PKR strongly inhibited replication of wild type VACV, which still contained both PKR inhibitors, whereas expression of Syrian or Armenian hamster PKR showed only strong reduction of either the K3L, or E3L deleted virus, respectively. Engineering PKR with enhanced resistance profiles to viral inhibitors might be a promising strategy to generate cells and organisms with broad resistance against viral PKR inhibitors and therefore increased resistance to viral infections.

It is noteworthy that VACV was able to replicate in Chinese hamster-derived V79-4 and Don cells. Previous reports have shown that VACV was unable to replicate in CHO cells, the most commonly used Chinese hamster cell line, whereas other orthopoxviruses, such as cowpox virus can replicate in these cells (55). The inability of VACV to replicate in CHO cells has been shown to be caused by a lack of a viral protein called CP77 (encoded by OPV023/CPXV025) that is found in other orthopoxviruses including monkeypox, taterapox and Akhmeta viruses (43). CP77 has been identified as an antagonist of the host restriction factor SAMD9L, and expression of Chinese hamster SAMD9L restricted VACV replication in human BT20 cells (56). Our finding that VACV was able to replicate in some Chinese hamster cells, indicates that the dependence of VACV on CP77 is due to cell line-specific differences rather than species-inherent differences.

The laboratory population of Syrian hamsters was severely bottlenecked from three litter mates (one male and two females) when it was established (57). Since our results show that Turkish hamster PKR was also resistant to E3 inhibition, the resistance of Syrian hamster PKR was thus not the result of this bottlenecking event. Because PKR from Armenian and Chinese hamsters were sensitive to E3 inhibition, the resistance of PKR from the two *Mesocricetus* species to E3 and the sensitivity of PKR from the more distantly related *Cricetulus* species indicate that resistance to an ancestral E3 (or similarly-acting viral dsRNA-binding protein) evolved sometime in an ancestor of both Syrian and Turkish hamster more than 2.5 - 2.7 million years ago (mya), but less than 7.6 - 10.8 mya when *Mesocricetus* and *Cricetulus* genera split (58).

E3 is an important poxvirus host range factor, which binds dsRNA. The most widely accepted model for E3-mediated inhibition of PKR, dsRNA-sequestration, is not species-specific; therefore it was unclear how E3 exerts its host range function. Here we describe a new species-specific mechanism that can explain this host range function. This discovery changes the paradigm for E3-host interactions and opens new research pathways for how E3 may interact with PKR to mediate species-specific inhibition during infection. The two most parsimonious explanations are either the formation of a heterodimer, in which E3 prevents PKR dimerization, or the formation of a heterotrimer, in which E3 disrupts productive dimerization of the kinase domains (Fig. 6). E3 resistance therefore could either be due to the exclusion of E3 from the homodimer or through structural differences, e.g. linker sequences, which would still allow PKR to form a functional kinase even if E3 is incorporated into the complex (Fig. 6). Taken together, this data shows a new pathway by which PKR and potentially other dsRNA binding host restriction factors may evolve resistance to viral antagonists.

**Fig 6.**
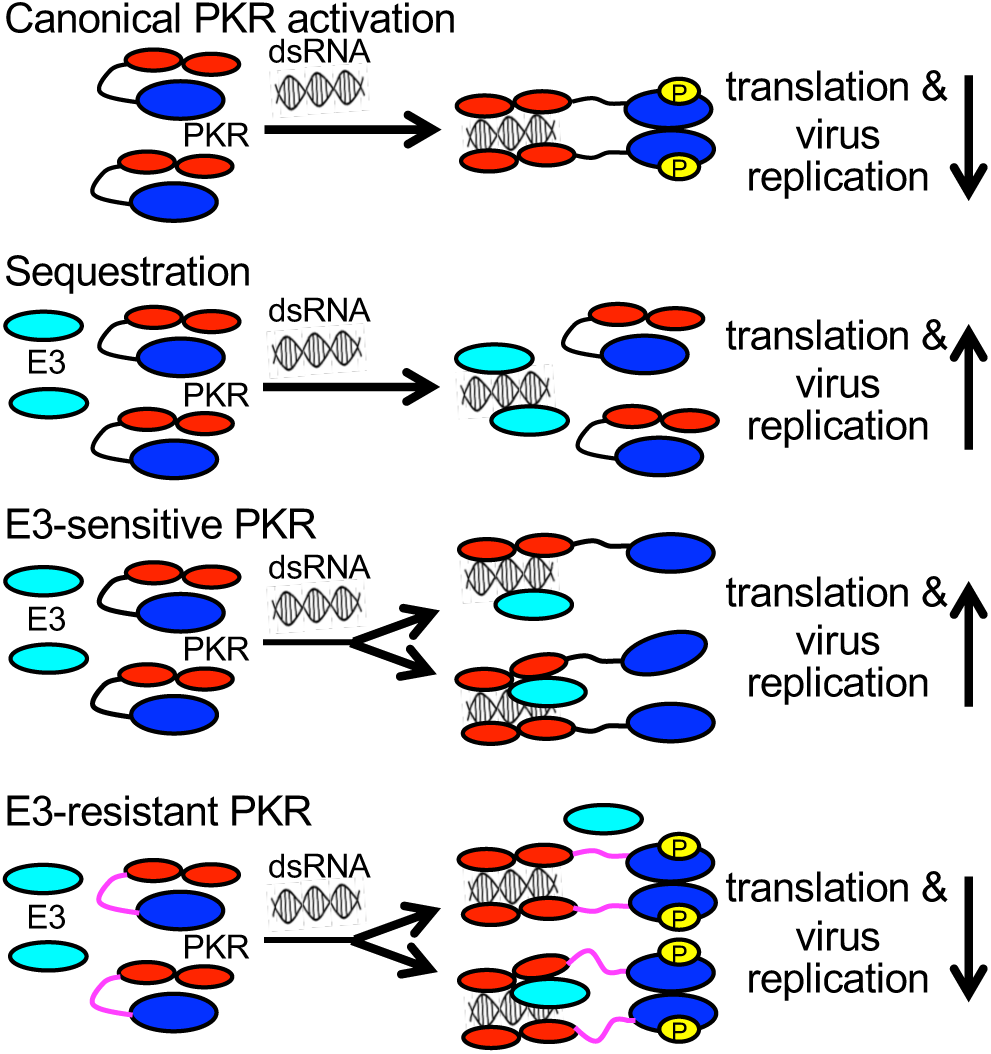
Models for PKR activation, and sensitivity to E3 inhibition. The effects of the outcomes on translation and virus replication are indicated by downwards (inhibition) or upwards (no inhibition) arrows. The following colors were used: red = dsRNA-binding domains of PKR; blue = PKR kinase domain; yellow = phosphorylation site in PKR; azure = E3.

## Materials and Methods

### Cell lines, viruses, yeast strains and plasmids

Syrian hamster BHK-21 (ATCC CCL-10), Armenian hamster AHL-1 (ATCC CCL-195), Chinese hamster Don (ATCC CCL-16), Chinese hamster V79-4 (ATCC CCL-93), human HeLa control and HeLa-PKR^kd^ (59) (both kindly provided by Dr. Charles Samuel) were maintained in Dulbecco’s Modified Essential Medium (DMEM, Life Technologies) supplemented with 5% fetal bovine serum (FBS, Fischer Scientific) and 25 µg/ml gentamycin (Quality Biologicals). RK13+E3L+K3L cells, which stably express VACV E3 and K3 (49), were grown in media supplemented with 500 µg/ml geneticin (Life Technologies) and 300 µg/ml zeocin (Life Technologies). Chinese hamster CHO-K1 (ATCC CCL-61; kindly provided by Dr. Anna Zolkiewska) cells were grown in RPMI medium (Life Technologies) supplemented with 10% FBS and 25 µg/ml gentamycin. Flp-In T-REx 293 cells (R78007, Fischer Scientific) were grown with 5% fetal bovine serum (FBS, Fischer Scientific) in Dulbecco’s Modified Essential Medium (DMEM, Life Technologies) supplemented with 100µg/mL zeocin (Life Technologies) and 15µg/mL blasticidin (Life Technologies). Flp-In T-REx 293 cell lines with reconstituted PKR were maintained in 5% FBS DMEM supplemented with 50µg/mL hygromycin B (Invitrogen) and 15µg/mL blasticidin. All cells were incubated at 37°C, 5% CO_2_.

### Knock-out of PKR in Flp-In T-REx 293 cells

PKR was knocked out in Flp-In T-REx 293 cells using the previously described guide RNA 5’-ATTCAGGACCTCCACATGAT-3’ (60), which targets the first coding PKR exon 3, and carried out using the Lipofectamine CRISPRMAX transfection reagent protocol (Fischer Scientific). HEK 293 Flp-In T-REx cells were plated in 24 well plates for a target confluency of ∼30%-40% for the following day. 24 hours later, gRNA, V2Cas9, OptiMEM, and CRISPRMAX reagents were diluted and added together following the manufacturer’s instructions. 50µL of the mix was then added to each well and the cells were left to incubate for 48 hours in a CO_2_ cell incubator. Cells were then washed with 500µL PBS and detached from the wells using 100µL trypsin. 1mL of fresh DMEM was added to each well after detachment of cells and subsequent cell clumps were broken apart by pipetting up and down on the bottom of the wells. The separated cells were then counted using a hemocytometer. A dilution of 0.15 cells per ml was then made using 10% DMEM 100µg zeocin/mL and 15µg/mL blasticidin. 200µL of the dilution was then added to each well of a 96 well plate. The cells were allowed to settle for 30 minutes and then each well was inspected and marked if one solitary cell was observed. From these single cells, clones for confirmation of the PKR KO by sequencing and VC-R4 infection were derived. The final Flp-In T-REx 293 PKR^ko^ cells were then named A2 for the well from which they were derived.

To analyze knock-out of PKR we extracted whole genomic DNA from candidate single cells clones, which were grown in 6 well plates to confluence, then trypsinized and washed with 1mL of PBS. Next cells were resuspended in 300µL of digestion buffer (100 mM NaCl,10mM Tris-Cl pH 8, 25mM EDTA pH 8, 0.5% SDS, 0.1mg/mL proteinase K) and proteinase K and incubated at 50°C for 2 hours and 30 minutes with gentle shaking (250 RPM) on a heat block. DNA was then extracted by adding 350µL phenol:chloroform:isoamyl, followed by centrifuging the samples at 1,700 RCF (Rotational Centrifugal Force) for 10 minutes and isolating the resulting aqueous layer. DNA was precipitated by adding 30µL 3M (molar) sodium acetate in 600µL 100% ethanol and pelleted at 1,700 RCF for 5 minutes. The pellet was then washed in 100µL 70% ethanol and centrifuged at 1,700 RCF for 5 minutes. Ethanol was discarded and the pellet allowed to dry before resuspension in 100µL ultrapure nuclease free H_2_O. We then ran PCR with the isolated gDNA and primers PKR-intron2-1F and PKR-intron3-1R (Table S3) surrounding human PKR exon 3, using the OneTaq polymerase (NEB). Following Topo TA Cloning protocol (Thermo Fisher Scientific), we combined PCR products with pCR4-TOPO plasmids and then transformed chemically competent *E. coli* via heat shock. We generated DNA mini-preps from 30 colonies with the expected inserts and sequenced them using Sanger sequencing with M13F (-21) and M13R. A single 17 bp deletion in exon 3 was observed in all 30 tested plasmid clones, and no intact PKR reads were detected.

For sequencing PKR from cDNA, 1.5 x 10^6^ Flp-In T-REx 293 PKR KO cells and T-REx™ 293 WT cells were seeded in six well plates. To induce transcription of PKR, the cells were infected with VC-R4 at an MOI of 3. At 6 hpi, the cells were lysed using TRI Reagent (Millipore Sigma, T9424). Extraction of total RNA and subsequent DNA digestion were performed according to Zymo Research RNA Extraction’s instruction. 1µg of total RNA was converted into cDNA with Protoscript II reverse transcriptase (NEB). The PKR open reading frame was amplified with primers PKR-exon2-F and PKR-exon17-R. PCR products were gel purified and Sanger sequenced. The identical 17 bp deletion in exon 3 was observed in all samples.

### Construction of PKR congenic Flp-In T-REx 293 cells

Flag-tagged Syrian hamster PKR (#793), Armenian hamster PKR (#794), Syrian hamster PKR I463D/I464F (#810), and human PKR (#836) were cloned into the pcDNA5/FRT expression vector, which confers hygromycin B resistance in successfully transfected cells. The resulting pcDNA5/FRT PKR-Flag plasmids were co-transfected into Flp-In T-REx 293 PKR^ko^ cells together with pOG44 plasmid, which expresses the Flp recombinase and recombines a Flp flanked gene of interest with the genomic Flp in site of Flp-In T-REx 293 cells. Transfection was carried out using SignaGen Genjet (SignaGen) following the manufacturer’s protocol. 4×10^6^ cells were transfected with 1µg of plasmids total (500ng each). Cells were then selected for successful plasmid uptake by adding 50µg/mL of Hygromycin B and cultivating surviving colonies. Total protein extracts were subjected to SDS page and western blot analysis using an anti-FLAG antibody with or without Doxycycline induction.

### Western blotting

Protein lysates were collected from confluent monolayers of cells grown in 6-well plates in 1% sodium dodecyl-sulfate (SDS) in PBS and sonicated 2×5 s to shear genomic DNA. Lysates from transfected cells were collected 48 hours after transfection in 1% SDS. Lysates from cells infected with wild-type VC-2 (Copenhagen), VC-R1 (ΔE3L), vP872 (ΔK3L), or VC-R2 (ΔE3LΔK3L) (MOI = 5) were collected at 6 hpi in 1% SDS for analyzing phosphorylation of eIF2α. All protein lysates were separated on 10% polyacrylamide gels and blotted on polyvinyl difluoride membranes (PVDF, GE Healthcare). Blotted membranes were blocked with either 5% non-fat milk or 5% BSA, before being incubated with rabbit anti-phospho-eIF2α (Ser51, BioSource International, kindly provided by Dr. Thomas Dever), rabbit anti-total eIF2α (Santa Cruz Biotechnology), or mouse anti-β-actin (Sigma-Aldrich) diluted in TBST buffer (20M Tris, 150mM NaCl, 0.1% Tween 20, pH 7.4). Membranes were incubated overnight at 4°C in the primary antibody, washed with TBST, and then incubated for one hour at room temperature with the secondary antibody conjugated to horseradish peroxidase (goat anti-rabbit-HRP or goat anti-mouse-HRP (Open Biosystems), donkey anti-rabbit-HRP or donkey anti-mouse-HRP (Life Technologies). The membranes were then washed with TBST to remove excess secondary antibody, and proteins were detected with Proto-Glo ECL detection buffers (National Diagnostics). Images were taken using a Kodak-4000MM Image Station or Invitrogen iBright Imaging. Mean band intensities for phosphorylated eIF2α to total eIF2α were quantified with ImageJ. The standard deviations of the ratios of phosphorylated eIF2α to total eIF2α for each sample were calculated from two independent experiments. FLAG-tagged PKR were detected with anti-FLAG M2 (Sigma-Aldrich) and goat anti-mouse (Invitrogen).

### Plasmids

Primers used in this study are shown in table S3. PKR from the indicated species and viral inhibitor genes were cloned into the pSG5 mammalian expression vector (Stratagene) for transient expression driven by the SV40 promoter as described (10). Each new construct was sequenced to confirm the absence of other mutations. The cloning of knock-down resistant human PKR, mouse PKR, Syrian hamster PKR, and Chinese hamster PKR into the pSG5 plasmid was described previously (34). Armenian hamster PKR (accession# MG702601) was cloned from AHL-1 cells using primers C42 x C40, located outside of the open reading frame (ORF) based on multiple sequence alignments from rodent PKRs and cloned into the pCR2.1-TOPO-TA vector (Invitrogen) for sequencing. Primers C47 and C48 were designed to sub-clone the PKR ORF into pSG5 with *SacI* and *XhoI* restriction sites. Turkish hamster PKR (accession# MG702602) was cloned from cDNA generated from total RNA isolated from the testes of a Turkish hamster (*Mesocricetus brandti*, kindly donated by Drs. Bob Johnston and Frank Castelli). Primers C42 x C40 were used to amplify the ORF of Turkish hamster PKR, and the primers used to sub-clone the Turkish hamster PKR ORF were BA70 and BA71. The E3L ortholog from VARV used is identical to that found in strain Somalia 1977 (accession# DQ437590, protein_id=ABG45218). Plasmids encoding for herpes Simplex virus-1 Us11 and mammalian reovirus σ3 (Type 1 Lang S4 gene), were kindly provided by Dr. Ian Mohr and Dr. Cathy Miller, respectively. For transfections, plasmids were prepared using the Nucleobond Xtra Midi Endotoxin Free DNA preparation kit (Macherey-Nagel).

### Yeast strains

The generation of the yeast strains stably transformed with empty vector (pRS305, J673), VARV E3L (J659), or VACV K3L-H47R (J674) at the LEU2 locus were described previously (10, 61). Syrian hamster PKR (pN1) and Armenian hamster PKR (pN2) were cloned into the pYX113 vector (R&D Systems), which encodes the GAL-CYC1 hybrid promoter and the selectable marker URA3. Yeast transformations using the lithium acetate/polyethylene glycol method and growth assays were carried out as previously described for human PKR (10). For each transformant, four single colonies were picked and three times colony purified. Purified transformants were streaked on SGal medium to induce PKR and inhibitor expression and grown at 30°C for 7 days.

### Phylogenetic and sequence analysis

Accession numbers of rodent PKR sequences are shown in table S1. Usage of designations of rodent families, subfamilies and genera is based on the classification used by the National Center for Biotechnology Information, taxonomy website (https://www.ncbi.nlm.nih.gov/taxonomy/). Protein sequence alignments and protein sequence identities of the PKRs were obtained using ClustalW in MegAlign (DNAStar, Inc.).

We reconstructed the phylogeny of rodent PKR sequences using the maximum likelihood approach (62), as implemented in PhyML (63). The input alignment included the European rabbit PKR as an outgroup and was generated using MUSCLE (64). In this phylogenetic analysis, we conducted 100 bootstrap iterations to estimate nodal support. We used FigTree v1.4.4 (http://tree.bio.ed.ac.uk/software/figtree/) to visualize the resulting phylogenetic tree.

In order to determine positively selected sites in hamster PKRs, we used the program codeml implemented in PAML (65). Residues with posterior probabilities greater than 95% in the model M8 were identified as sites under positive selection (μ > 1) in BEB (Bayes Empirical Bayes) analysis (66).

### Virus infections, interferon and siRNA treatments

Viruses used in this study are VC-2 (VACV-Copenhagen) and its derivatives vP872, whose K3L gene is deleted (ΔK3L) (27) (both kindly provided by Dr. Bert Jacobs), VC-R2 (ΔE3L, ΔK3L), a derivative of vP872, in which E3L was replaced by a destabilized EGFP gene (67) and VC-R1 (ΔE3L) which was generated by replacing E3L with a destabilized EGFP gene in VC-2 as described for VC-R2.

For all virus infections, sonicated virus samples were diluted in DMEM (or RPMI) supplemented with 2.5% FBS to perform infections at the indicated multiplicities of infection (MOI). For each of the cell lines used, 5.0 x 10^5^ cells were plated in 6-well plates one day before infection, and infections were performed in duplicate unless otherwise noted. The growth media was removed from each well before adding the diluted virus inoculum and incubating it for one hour at 37°C. After the incubation period, the inoculum was removed, cells were washed twice with phosphate buffered saline (PBS), and fresh growth media was added. Virus was collected at the indicated times post infection by scraping cells directly into the media and lysates were subjected to three rounds of freeze/thaw cycles followed by sonication (2 x 15 s) in a cup horn sonicator (QSonica). Virus titers were measured in plaque forming units per ml (pfu/ml) on RK13+E3L+K3L cells.

To investigate induction of PKR expression, cells were pre-treated with 5 units/ml mouse interferon-α1 (mmIFNα 1, pbl interferon source) 17 hours prior to collecting RNA for RT-PCR analysis. To analyze the effect of Interferon treatment on virus replication, BHK cells were pre-treated with 50 units/ml mmIFNα1, 24 hours prior to infection. Four siRNA duplexes (21nt) were designed and synthesized using the Dharmacon siDESIGN Center (dharmacon.gelifesciences.com) to target Syrian hamster PKR and Armenian hamster PKR (Dharmacon, Table S4). Cells were transfected with the pooled siRNA duplexes or mock-transfected in 6-well plates with 50nM diluted in siRNA buffer (60mM KCl, 6mM HEPES, 20µM MgCl_2_, pH 7.4) using Lipofectamine RNAiMAX siRNA transfection reagent (Invitrogen) according to the manufacturers’ instructions. The media in each well was changed after 24 hours, and the cells were infected with the four VACV strains at MOI = 0.01, 24 hours after transfection. Virus lysates from two replicate infections were collected after 30 hours and titered on RK13+E3L+K3L cells.

### RT-PCR

RNA was collected from mmIFNα1 treated or untreated cells grown in 6-well plates 17 hours after treatment using TRIzol (Thermo Fisher Scientific). cDNA was then generated using SuperScript III reverse transcriptase and oligo-dT primers (Invitrogen). Primers used to amplify Syrian hamster PKR from BHK-21 cells for RT-PCR analysis were BA16 and BA26. Primers used to amplify Armenian hamster PKR from AHL-1 cells were BA14 and BA27. Primers used to amplify Chinese hamster PKR from V79-4 cells were C49 and C50. Primers used to amplify eIF2a from all cell lines were eIF2a-1F and eIF2a-2R.

### Luciferase-based reporter assay for PKR inhibition

The luciferase-based reporter assay for inhibition of PKR activity was described previously (10). Briefly, 5 x 10^4^ HeLa-PKR^kd^ cells were seeded 24 hours before transfection in 24-well plates. For each transfection, 0.05µg of firefly luciferase encoding plasmid (pGL3promoter, Promega), 0.2 µg PKR encoding plasmids (pSG5), and 0.4 µg VACV E3L or VACV K3L were transfected using GenJet-Hela (SignaGen) in triplicates. For titration experiments, VACV E3L was co-transfected at the indicated concentrations and the total amount of plasmid transfected was kept constant with additional empty vector (pSG5). Cell lysates were harvested 48 hours after transfection using mammalian lysis buffer (Goldbio), and the luciferase activity was determined by measuring luminescence in a luminometer (Berthold) after adding luciferin substrate (Promega). Luciferase activity from vector control transfections were compared to transfections with only PKR encoding plasmids to assess the PKR activity for each species, which was then used to normalize co-transfections of the corresponding PKR with each viral inhibitor.

## Acknowledgments.

We are grateful to Drs. Thomas Dever and Bernard Moss for their support and advice in the early stages of this project. We thank Drs. Bertram Jacobs, Charles Samuel, Anna Zolkiewska, Bob Johnston, Frank Castelli, Ian Mohr, Cathy Miller and Thomas Dever for generously providing materials and reagents. Parts of this work were included in the PhD dissertation of S. L. H. (68). This work was supported by Grant R01 AI114851 (to S.R.) from the National Institute of Allergy and Infectious Diseases.

