## Supporting information for "Molecular basis for the host range function of the poxvirus PKR inhibitor E3"

**A**

```
                                                dsRBD1
Ma PKR  MASDTPGFYMDKLNKYHQKHRVTITYKHLFTTGPPHDRRFTFQVIIDGEEFPEAEGKTKQEAKNAAA  67
Mb PKR  MASDTPGFYMDKLNKYHQKHRVTITYKHLFTTGPPHDRRFTFQVIIDGEKFPEAEGKTKQEAKNAAA  67
Cm PKR  MASDTPGFYMDKLNKYHQKHKVKITYKQLHITGPPHDRRFTFQVIIDGEEFPEAEGRTKQEAKNAAA  67
Cg PKR  MASDTPGFYMDKLNKYHQKHKVMITYKQLHITGPPHDRRFTFQVIINGEEFPEAEGRTKQEAKNAAA  67
        dsRBD1                                          dsRBD2
Ma PKR  KLAVDTLNANKANSHTDALEGSFTGNYIGLVNSYAQKEKLSVNYQQCAFNTQSPQRFCYKCIIGLKT  134
Mb PKR  KLAVDKLNANKANSHTDALEGSFTGNYIGLVNSYAQKEKLSVNYQQCAFNTQSPQRFCYKCIIGLKT  134
Cm PKR  QLAVERLNENKVDSHTDTSEGSLSGNYIGLVNSYAQKENLSVNYEQCTSNTQSPQRFCCKCKIGPKT  134
Cg PKR  QLAVERLNENKADSQTDASEGLLSGNYIGLVNSYAQKENLSVNYEQCTSNTQSPQRFCCKCKIGLKT  134
                                                    sv1
Ma PKR  YGIGSGATKQEAKQLAAKEAYQKLPEKSSMRVCRASSGLSKSSSSGHFNSWSTANNFTSLSAPGSDF  201
Mb PKR  YGIGSGATKQEAKQLAAKEAYQKLPEKSSMRVCRASSGLSKSSSSGHFNSWSTANKFTSLSAPGSDV  201
Cm PKR  YGIGSGATKQEAKQLAAKDAYQKLSEKSSMRVDRASSDLSTSSSSGRSSSLCTTSNFSSLSAPESDF  201
Cg PKR  YGIGSGATKQEAKQSAAKDAYQKLSEKSSMRVDRTSSGLSTSSSSGLSSSLSTASHFTSLSAPQSDF  201
            sv1          R2    R3     R4    R5
Ma PKR  SETASNNHC--VFFKSPLMNGLRENKRKPRVKLSSNDVQRNKYTTDSRFNEDFEDIEEIGSGGFGQV  266
Mb PKR  SATASNNHC--VFFKSPLMNGLRENKRKPPVKLSSNDVQSNKYTTDSRFNEDFEDIEEIGSGGFGQV  266
Cm PKR  SETASQNHCHSVSFESPFTNGLRENKRKSGVKPLLNDVQRSKYTIDPRFNDDFEDIEEIGSGGFGQV  268
Cg PKR  SEIASKNHWHSISFESPFTNGLRENKRKSGVKPSPSDVQRNKYTMDSRFNNDFEDIEEIGSGGFGQV  268
                                 KD                               KI
Ma PKR  FKAKHRIDGKTYAIKRVKYNSEKEVREVKALAALSHVNIVQYHFCWLGQDWEYDTEQSMNSATPRSK  333
Mb PKR  FKAKHRIDGKTYAIKRVKYNSEKEVREVKALAALSHVNIVQYHFCWLGQDWEYDTEQSMNSATPRSK  333
Cm PKR  FKAKHRIDGKTYAIKRVRYNTEKEVREVKALAALSHVNIVQYHSCWLGQD--YAPEHSLDRNPRRAK  333
Cg PKR  FKAKHRIDGKTYAIKRVSCNTEKEVREVKALAALSHINIVQYHSCWLGQD--YDPEHSPDRNPRRSK  333
                                    KD
Ma PKR  TECLFIQMEFCDKGTLEQWMTSVKQRKVDNDLFLELAEQITTGVDYIHSKGLIHRDLKPSNIFLVDE  400
Mb PKR  TECLFIQMEFCDEGTLEQWMTSVKQRKVDNDLFLELAEQITTGVDYIHSKGLIHRDLKPSNIFLVDE  400
Cm PKR  TDCLFIQMEFYDKGTLEQWMLNCKQRKVDNDLFLEFSEQITTGVDYMHSKGLIHRDLKPCNIFLVDE  400
Cg PKR  TQCLFIQMEFYDKGTLEQWMLNCKQRKVDNDLFLELSEQITTGVDYIHSKGLIHRDLKPCNIFLVDE  400

Ma PKR  KHIKIGDFGLVTALENDGNRTKNTGSLLYMSPEQLSLQEYGKEVDIFALGLILAELLHICITIIEKS  467
Mb PKR  KHIKIGDFGLVTALENDGNRTKNTGSLLYMSPEQLSLQEYGKEVDIFALGLILAELLHICITIVEKS  467
Cm PKR  KHIKIGDFGLVASLEKDGNRTKGTGTELYMSPEQLSLQEYGKEVDIFALGLILAELLHICITDFEKI  467
Cg PKR  KHIKIGDFGLVTSMEKDGNRTKRTGTLLYMSPEQLSLQEYGKEVDIFALGLILGELLHICITDLEKS  467

Ma PKR  KFFTDLRDGIFHDDIFGSKEKRLLTKLLSKKPTDRPSASEILMTLAEWKNISEVQKRHTH  527
Mb PKR  KFFTDLRDGIFHDDIFGSKEKRLLMKLLSKKPTDRPSAHEILKTLAEWKSISEVQKRYTH  527
Cm PKR  KFFSDLRAGIFHDNIFGSKEKNLLMKLLSNKPTERPSTSEILMTLAEWKRISEIQKRHTR  527
Cg PKR  KFFNDLRDGIFHDDIFGSKEKSLLKKLLAKKPMERPSTSEILMTLAEWKSISEIQKRHTL  527
```

**B**

| | Ma | Mb | Cm | Cg | Mm | Hs |
|---|---|---|---|---|---|---|
| Ma (Syrian hamster) PKR | ------ | 97.3 | 82.1 | 83 | 71.3 | 62.8 |
| Mb (Turkish hamster) PKR | 97.3 | ------ | 80.6 | 81.7 | 70.3 | 61.8 |
| Cm (Armenian hamster) PKR | 82.1 | 80.6 | ------ | 89.9 | 71.1 | 61.3 |
| Cg (Chinese hamster) PKR | 83 | 81.7 | 89.9 | ------ | 71.9 | 62.4 |
| Mm (mouse) PKR | 71.3 | 70.3 | 71.1 | 71.9 | ------ | 61.1 |
| Hs (human) PKR | 62.8 | 61.8 | 61.3 | 62.4 | 61.1 | ------ |

**Fig. S1. Multiple sequence alignment of PKR from different hamster species.** (A) The amino acid sequences of Syrian hamster (Ma) PKR, Turkish hamster (Mb) PKR, Armenian hamster (Cm) PKR, and Chinese hamster (Cg) PKR were aligned by Clustal W (MegAlign, DNAStar, Inc.). The positions of the two dsRNA-binding domains (dsRBD), kinase domain (KD) and the kinase insert domain (KI) are indicated above the alignment. Residues differing from Syrian hamster PKR are shown in bold. Residues in the Cricetulus (Cm and Cg) PKRs that differ from residues in Mesocricetus (Ma and Mb) PKRs are highlighted in red. Differences between Cm and Cg PKR are highlighted in blue. Residues missing in the Syrian hamster PKR splice variant (sv1) and additional removed blocks (R2-R5) in deletion constructs in the linker are boxed. (B) The amino acid sequence identities between the four hamsters, mouse and human PKR as calculated from a multiple sequence alignment using MegAlign are shown.

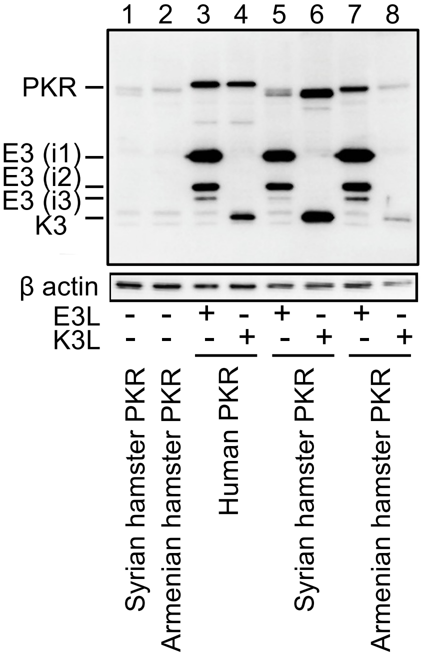


**Fig. S2. Expression of PKR, E3, and K3 in transfected cells**. HeLa-PKR$^{kd}$ cells were transfected with plasmids encoding FLAG-tagged PKR from the indicated species (1 µg) with or without FLAG-tagged VACV K3L or E3L (1 µg). 1% SDS protein lysates were collected and analyzed by western blot for gene expression 24 hours after transfection with anti-FLAG antibodies (top panels) or anti-*β* actin as a loading control (bottom panels). E3 isoforms, likely generated by alternative start-codon usage, are labeled i1, i2 and i3.

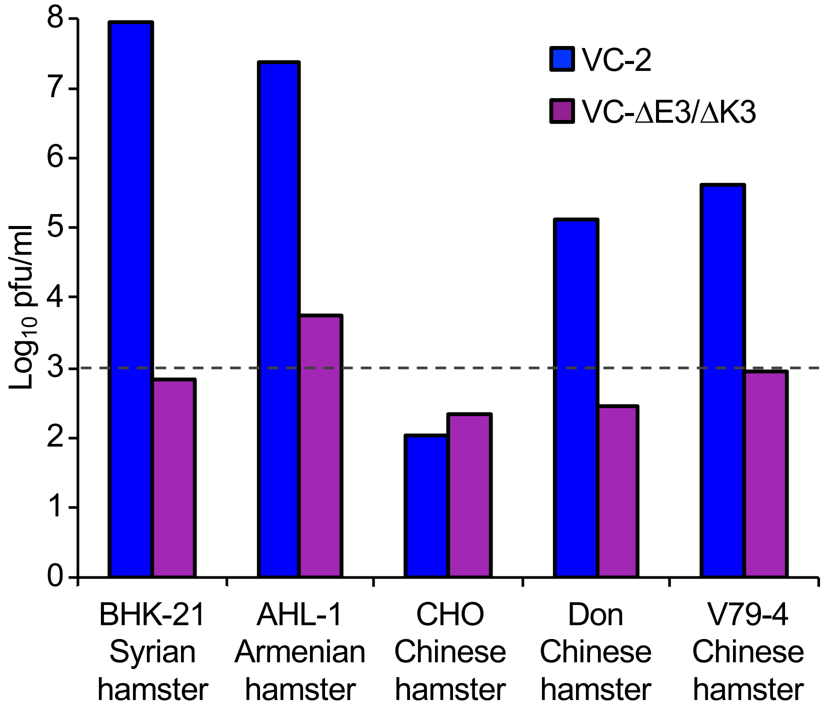


**Fig. S3. VACV replication in cells from different hamster species.** Cells derived from Syrian hamster (BHK-21), Armenian hamster (AHL-1), and Chinese hamsters (CHO, Don, V79-4) were infected with wild-type VACV-Cop (VC-2) or VC-R2 lacking E3L and K3L at MOI = 0.001. Virus was collected after 48 hours and titered on RK13+E3L+K3L cells. The dashed line represents the level of input virus.

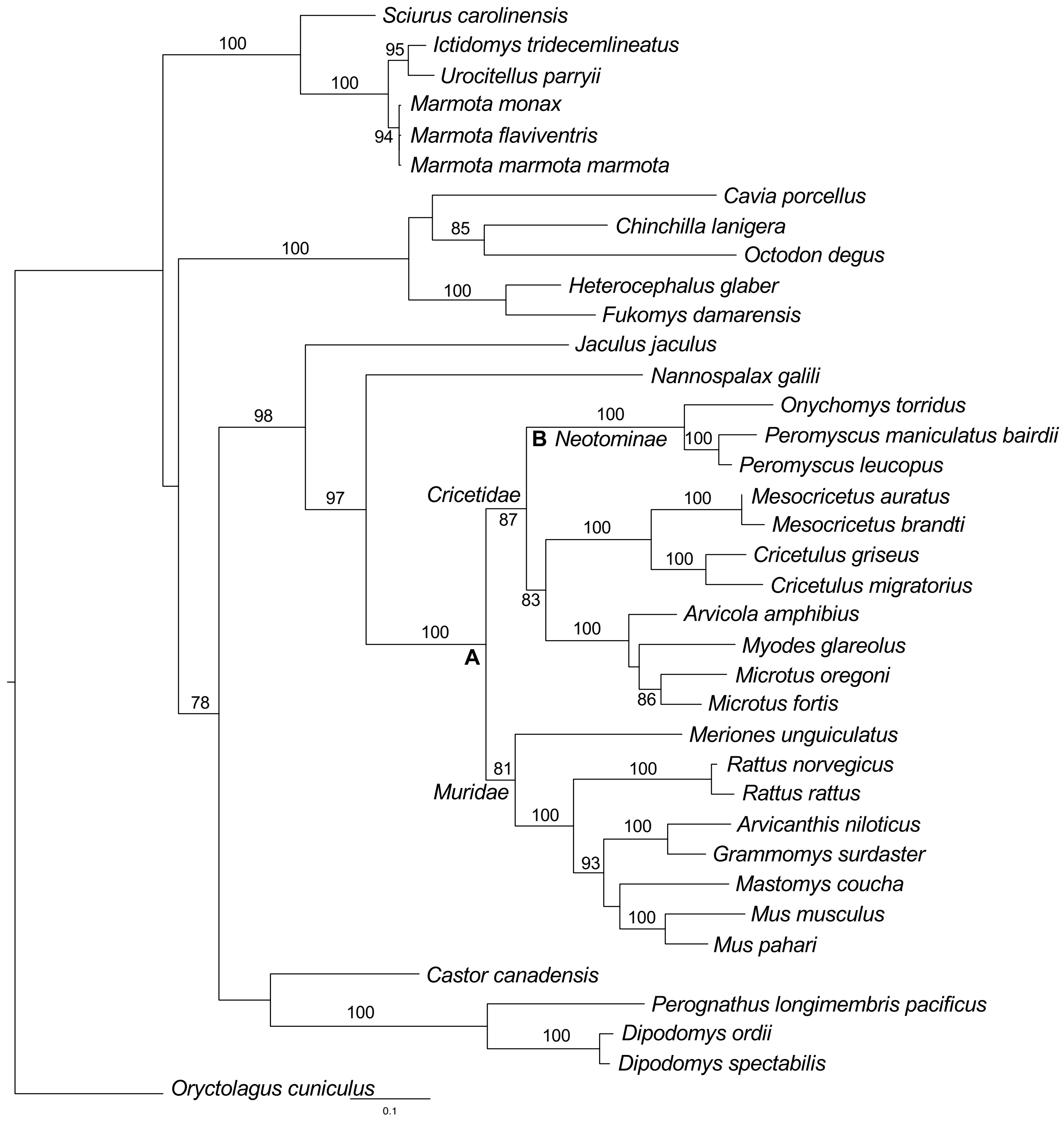


**Fig. S4. Phylogenetic analysis of rodent PKR sequences.** Phylogenetic tree was constructed with a multiple sequence alignment of 36 rodent PKR sequences and European rabbit (*O. cuniculus*) PKR (as outgroup), generated with MUSCLE using maximum likelihood analysis (PhyML). Bootstrap support (> 70) form 100 replicates is shown on nodes. The tree was rooted to *O. cuniculus* PKR.

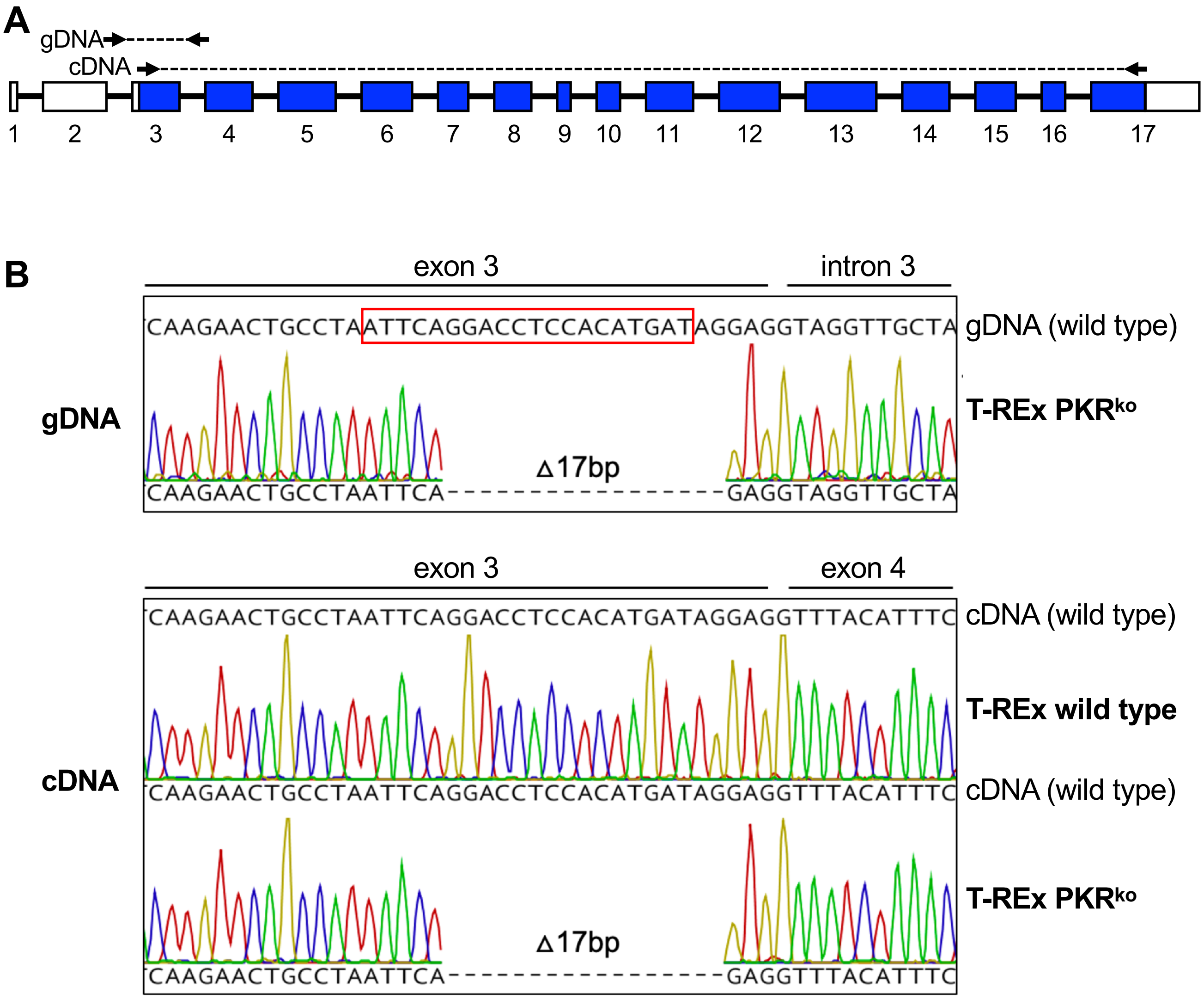


**Fig. S5. A 17bp deletion in exon 3 of PKR in T-REx PKR$^{ko}$ cells disrupts the open reading frame.** (A) The exon/intron structure of the human PKR gene is shown. The open reading frame (ORF) is highlighted in blue. Genomic DNA and total RNA was extracted from T-REx PKR$^{ko}$ cells. From genomic (g) DNA, PCR was performed with primers in intron 2 and intron 3 as indicated by the arrows. From cDNA, PCR was performed with primers in exon 3 and exon 17, as indicated. (B) A red box highlights the sequence that corresponds to the guide RNA used for CRISPR/Cas9-mediated knock-out. After Topo TA Cloning of the PCR products from gDNA, plasmids from 30 colonies were Sanger sequenced. All sequences contained an identical 17 bp deletion that disrupts the open reading frame. PCR using cDNA, yielded single bands from wild type T-REx cells and T-REx PKR$^{ko}$ cells, which were directly sequenced after gel purification. Both PCR products yielded clean chromatograms, with that obtained from wild type T-REx cells showing an uninterrupted ORF, whereas the identical 17 bp deletion as obtained from gDNA was identified from T-REx PKR$^{ko}$ cells.

**Table S1. Genes included in phylogenetic analysis**

| Species | Common name | Accession number | Family |
|---|---|---|---|
| *Mesocricetus auratus (Ma)* | Syrian hamster | NM_001281945.1 | *Cricetidae* |
| *Mesocricetus brandti (Mb)* | Turkish hamster | MG702602.1 | *Cricetidae* |
| *Cricetulus griseus (Cg)* | Chinese hamster | KT272869.1 | *Cricetidae* |
| *Cricetulus migratorius (Cm)* | Armenian hamster | MG702601.1 | *Cricetidae* |
| *Arvicola amphibius (Aa)* | Eurasian water vole | XM_038320475.1 | *Cricetidae* |
| *Microtus fortis (Mf)* | reed vole | XM_050163535 | *Cricetidae* |
| *Microtus oregoni (Mo)* | creeping vole | XM_041631130 | *Cricetidae* |
| *Myodes glareolus (Mg)* | Bank vole | XM_048416055 | *Cricetidae* |
| *Onychomys torridus (Ot)* | southern grasshopper mouse | XM_036170876.1 | *Cricetidae* |
| *Peromyscus leucopus (Pl)* | white-footed mouse | XM_028873276.2 | *Cricetidae* |
| *Peromyscus maniculatus bairdii (Pm)* | prairie deer mouse | XM_042266732.1 | *Cricetidae* |
| *Mastomys coucha (Mc)* | Southern multimammate mouse | XM_031356301.1 | *Muridae* |
| *Mus pahari (Mp)* | shrew mouse | XM_021218187.2 | *Muridae* |
| *Meriones unguiculatus (Mg)* | Mongolian gerbil | XM_021661519.1 | *Muridae* |
| *Arvicanthis niloticus (Nile rat)* | Nile rat | XM_034514534.1 | *Muridae* |
| *Grammomys surdaster (Gs)* | African woodland thicket rat | XM_028773442.1 | *Muridae* |
| *Mus musculus (Mm)* | house mouse | BC016422.1 | *Muridae* |
| *Rattus norvegicus (Rn)* | Norway rat | L29281.1 | *Muridae* |
| *Rattus rattus (rr)* | black rat | XM_032908889.1 | *Muridae* |
| *Nannospalax galili (Ng)* | Upper Galilee mountains blind mole rat | XM_008822598.3 | *Spalacidae* |
| *Jaculus jaculus (Jj)* | lesser Egyptian jerboa | XM_045137788.1 | *Dipodidae* |
| *Heterocephalus glaber (Hg)* | naked mole-rat | XM_004839297.3 | *Bathyergidae* |
| *Fukomys damarensis (Fd)* | Damara mole-rat | XM_010611405.3 | *Bathyergidae* |
| *Octodon degus (Od)* | degu | XM_023711201.1 | *Octodontidae* |
| *Chinchilla lanigera (Cl)* | long-tailed chinchilla | XM_013517630.1 | *Chinchillidae* |
| *Cavia porcellus (Cp)* | Guinea pig | KT272870.1 | *Caviidae* |
| *Sciurus carolinensis (Sc)* | gray squirrel | XM_047523348.1 | *Sciuridae* |
| *Marmota marmota marmota (Mmm)* | Alpine marmot | XM_015479767.2 | *Sciuridae* |
| *Marmota monax (Mmo)* | woodchuck | XM_046441648.1 | *Sciuridae* |
| *Marmota flaviventris (Mf)* | yellow-bellied marmot | XM_027933411.2 | *Sciuridae* |
| *Ictidomys tridecemlineatus (It)* | thirteen-lined ground squirrel | XM_005336548.4 | *Sciuridae* |
| *Urocitellus parryii (Up)* | Arctic ground squirrel | XM_026405873.1 | *Sciuridae* |
| *Castor canadensis (Cc)* | American beaver | XM_020167142.1 | *Castoridae* |
| *Perognathus longimembris pacificus (Plp)* | Pacific pocket mouse | XM_048353124.1 | *Heteromyidae* |
| *Dipodomys spectabilis (Ds)* | banner-tailed kangaroo rat | XM_042675531.1 | *Heteromyidae* |
| *Dipodomys ordii (Do)* | Ord's kangaroo rat | XM_013013683.1 | *Heteromyidae* |
| *Oryctolagus cuniculus (Oc)* | European rabbit | KT272867.1 | *Leporidae* |

**Table S2. Positively selected sites in rodent PKRs.**

| Residue (S. hamster) | p values Clade A *Cricetidae* + *Muridae* | p values Clade A - clade B *Cricetidae* + *Muridae* - *Neotominae* |
|---|---|---|
| F30 | 0.990** | 0.980* |
| T91 | 0.932 | 0.966* |
| S120 | 0.976* | 0.928 |
| C125 | 0.960* | 0.993** |
| H181 | 0.998** | 0.998** |
| C210 | 0.991** | 0.916 |
| P216 | 0.965* | 0.973* |
| S246 | 0.991** | 0.983* |
| E250 | 0.968* | 0.992** |
| F310 | 0.985* | 0.828 |
| Q323 | 0.941 | 0.977* |
| M325 | 0.980* | 0.977* |
| E335 | 0.981* | 0.975* |
| F366 | 0.962* | 0.955* |
| N423 | 1.000** | 1.000** |
| L427 | 0.995** | 0.952* |
| Q438 | 0.999** | 0.990** |
| I461 | 0.997** | 0.986* |
| I463 | 0.980* | 0.952* |
| I464 | 0.999** | 0.998** |
| S467 | 0.988* | 0.988* |
| T471 | 0.933 | 0.951* |

** indicate p values ≥ 0.99; * indicate p values ≥ 0.95

**Table S3. Primers used in this study**

| Primer name | Sequence (5' -> 3') |
|---|---|
| C42 | GGG CGA CGC GAT CTC AGA GTC AGC ACC CGA AGC AAA AGT CGA ATC CT |
| C40 | GGA AAA AAA AGT ACA ATG TTC CCC CTT ATT CCA TCT CAG ATT TTA G |
| C47 | TAA GAG CTC GCC ACC ATG GCC AGT GAT ACA CCG GG |
| C48 | AAT CTC GAG TCA CTA ACG TGT GTG TCT TTT CTG TAT C |
| BA70 | GTA CGA GCT CGC CAC CAT GGC CAG TGA TAC ACC C |
| BA71 | CTG TCT CGA GTC ACT AAT GTG TGT ATC GTT TCT GTA CTT CTG |
| BA16 | TCG CTA GCA TGG CCA GTG ATA CTC CC |
| BA26 | TAA TCT CGA GAT GTG TGT GTC GTT TCT GTA CTT C |
| BA14 | ACT GCT AGC ATG GCC AGT GAT ACA CCG G |
| BA27 | TAA TCT CGA GAC GTG TGT GTC TTT TCT GTA TCT C |
| C49 | TAA GAG CTC GCC ACC ATG GCC AGT GAT ACA CCG GGT TTC TAC ATG GAC |
| C50 | AAT CTC GAG TCA CTA AAG TGT GTG TCT TTT CTG TAT CTC |
| eIF2a-1F | GTA GTG ATG GTG AAT GTA AGA TCC |
| eIF2a-2R | CAT CAC ATA CCT GGG TGG AG |
| PKR-intron2-F | TTG TAA AAC GAC GGC CAG TGA CCC CTC TGT CTC CTA AA |
| PKR-intron3-R | ATC CCC GGG TAC CGA GCT CGC CTA TGA GTG AGA ACA TGC |
| PKR-exon2-F | TCC AGC ACA GTG GCG GCC ACC ATG GCT GGT GATC |
| PKR-exon17-R | TTT AAA CGG GCC CTC TAG ACC TAC TAA CAT GTG TGT CGT TCA TTT TTC TC |

**Table S4. siRNA duplexes used in this study**

| siRNA duplexes | Sequence (5' -> 3') |
|---|---|
| duplex 1 | GGA AUU AGC UGA ACA AAU AUU, UAU UUG UUC AGC UAA UUC CUU |
| duplex 2 | CAC CAG AAC GAU AGA GUA AUU, UUA CUC UAU GCU UCU GGU GUU |
| duplex 3 | CCA CAU GAC AGA AGG UUU AUU, UAA ACC UUC UGU CAU GUG GUU |
| duplex 4 | GGA AAG UAG ACA AUG AUU UUU, AAA UCA UUG UCU ACU UUC CUU |
